# A trade-off between resistance to infection and reproduction in primate evolution

**DOI:** 10.1101/2020.07.10.186742

**Authors:** Sumnima Singh, Jessica A. Thompson, Sebastian Weis, Daniel Sobral, Mauro Truglio, Bahtiyar Yilmaz, Sofia Rebelo, Silvia Cardoso, Erida Gjini, Gabriel Nuñez, Miguel P. Soares

## Abstract

Most mammals express a functional *GGTA1* gene encoding the N-acetyllactosaminide α-1,3-galactosyltransferase enzyme, which synthesizes Galα1-3Galβ1-4GlcNAc (αGal) and are thus tolerant to this self-expressed glycan epitope. Old World primates including humans, however, carry *GGTA1* loss-of-function mutations and lack αGal. Presumably, fixation of such mutations was propelled by natural selection, favoring the emergence of αGal-specific immunity, which conferred resistance to αGal-expressing pathogens. Here we show that loss of *Ggta1* function in mice enhances resistance to bacterial sepsis, irrespectively of αGal-specific immunity. Rather, the absence of αGal from IgG-associated glycans increases IgG effector function, via a mechanism associated with enhanced IgG-Fc gamma receptor (FcγR) binding. The ensuing survival advantage against sepsis comes alongside a cost of earlier onset of reproductive senescence. Mathematical modeling of this trade-off shows that under conditions of high exposure to virulent pathogens, selective pressure can fix *GGTA1* loss-of-function mutations, as likely occurred during the evolution of primates towards humans.

## INTRODUCTION

Mammals, including humans, generate relatively high levels of circulating anti-glycan antibodies (Ab) at steady state (Huflejt et al., 2009; Kearney et al., 2015; Schneider et al., 2015; Stowell et al., 2014). These are referred to as natural antibodies (NAb), to hint that their generation occurs in the absence of traceable immunization. It is becoming clear, however, that circulating anti-glycan NAb are generated to a large extent in response to glycans expressed by immunogenic bacteria (Palm et al., 2014) in the gut microbiota (Bunker et al., 2017; Soares and Yilmaz, 2016; Yilmaz et al., 2014).

Circulating NAb of the IgM isotype provide an immediate lytic response to pathogens via activation of the complement-cascade (Ochsenbein et al., 1999; Yilmaz et al., 2014). In contrast, circulating NAb from the IgG isotype confer protection against infection via cognate interaction with Fcγ receptors (FcγR) (Zeng et al., 2016) expressed by immune cells, a central driver of IgG effector function (Lu et al., 2018; Ravetch and Kinet, 1991).

The protective effect exerted by glycan-specific NAb is operational only when the targeted glycans are not expressed as self epitopes (Soares and Yilmaz, 2016). Remarkably, this fundamental principle of immune self/non-self discrimination was circumvented for several glycans, through the natural selection and fixation of loss-of-function mutations in genes synthesizing those glycans (Soares and Yilmaz, 2016; Springer and Gagneux, 2016). This is perhaps best illustrated for the loss of αGal expression in ancestral Old World primates, including humans, as originally described by K. Landsteiner and P. Miller (Galili et al., 1987; Galili and Swanson, 1991; Landsteiner and Miller, 1925).

The αGal glycan is synthesized in most mammals, including New World monkeys, by GGTA1, a galactosyltransferase that catalyzes the transfer of a galactose (Gal) in α1-3 linkage, from a uridyl-diphosphate (UDP) donor onto the N-acetyllactosamine (Galβ1,4GlcNAc-R) of glycoproteins. This reaction does not occur in Old World primates, including humans (Galili et al., 1988b), which carry a *GGTA1* pseudogene (Galili et al., 1987; Galili and Swanson, 1991). Presumably, loss of *GGTA1* function in the ancestor of Old World primates contributed to shaping the human anti-glycan NAb repertoire (Huflejt et al., 2009), allowing for the emergence of αGal-specific immunity (Galili et al., 1984; Macher and Galili, 2008; Soares and Yilmaz, 2016).

When considering infection as a major driving force in evolution (Haldane, 1949), it is reasonable to assume that protection from infection should act as a major driving force in the natural selection and fixation of *GGTA1* loss-of-function mutations during primate evolution. In strong support of this notion, circulating αGal-specific NAb can account for up to 1-5% of all circulating IgM and IgG in healthy adult humans (Macher and Galili, 2008), providing protection against infection by pathogens expressing αGal-like glycans (Soares and Yilmaz, 2016; Takeuchi et al., 1996; Yilmaz et al., 2014).

Sepsis is a life-threatening organ dysfunction caused by a deregulated host response to infection (Singer et al., 2016), which can account for as much as 20% of global human mortality (Rudd et al., 2020). Presumably, therefore, infections by virulent pathogens that can trigger the development of sepsis are likely to have exerted a major selective pressure throughout primate and human evolution. Sepsis is thought to originate and progress, to at least some extent, from the immune response elicited upon translocation across epithelial barriers of bacterial pathobionts from the microbiota (Rudd et al., 2020; Vincent et al., 2009). As circulating αGal-specific NAb are generated, at steady state, in response to αGal-like glycans expressed by immunogenic bacteria in the microbiota (Galili et al., 1988a; Soares and Yilmaz, 2016; Springer and Horton, 1969), one might expect αGal-specific NAb to provide some level of protection against sepsis. While there is no epidemiological evidence to support this notion, there are other mechanisms via which loss of *GGTA1* function could enhance protection against bacterial sepsis.

Circulating IgG carry bi-antennary glycan structures, N-linked to an evolutionarily conserved asparagine-297 in their constant heavy chain of the Fc domain (Anthony et al., 2012). These glycan structures are composed of varying amounts of N-acetylglucosamine (GlcNAc), fucose, mannose, galactose (Gal) and sialic acid molecules, which modulate the binding affinity of the IgG Fc domain to different FcγR and proteins of the complement cascade (Anthony et al., 2012; Dekkers et al., 2017; Wang and Ravetch, 2019). Given that αGal is part of these IgG-associated glycan structures in mammals that carry a functional *GGTA1* gene (de Haan et al., 2017), we hypothesized that αGal can modulate IgG binding to FcγR and/or complement proteins (Anthony et al., 2012; Dekkers et al., 2017; Wang and Ravetch, 2019). In support of this hypothesis, when present in IgG-associated glycan structures terminal Gal residues can modulate IgG-FcγR binding and complement activation (Nimmerjahn et al., 2007). This is also the case for other glycan residues, as demonstrated for example for fucose in an α1,6-linkage (Nimmerjahn and Ravetch, 2005).

Here we hypothesized that the presence or absence of αGal from the glycan structure of IgG might modulate IgG NAb effector function in a manner that modulates resistance to bacterial sepsis. We report that loss of *Ggta1* function in mice confers robust protection from bacterial sepsis via a mechanism that acts irrespectively of αGal-specific immunity. This robust protective effect is mediated, instead, by enhancement of IgG NAb effector function, via a mechanism associated with increased IgG-FcγR binding. The gained fitness advantage against infection is associated, however, with an accelerated onset of reproductive senescence. Mathematical modeling of this evolutionary trade-off suggests that under conditions of high exposure to virulent pathogens, the fitness gain prevails over the cost, providing a possible explanation for the natural selection and fixation of *GGTA1* loss-of-function mutations during primate evolution towards humans.

## Results

### Loss of *Ggta1* function in mice enhances resistance to systemic infection by symbiotic gut bacteria

We tested the hypothesis that the loss of *GGTA1* function during primate evolution (Galili and Swanson, 1991) might provide protection against bacterial sepsis in *Ggta1*-deficient (*Ggta1*^*-/-*^) mice, a well-established experimental model that mimics the absence of *GGTA1* function in modern humans. Polymicrobial infection was induced by cecal ligation and puncture (CLP), a well-established model of sepsis (Rittirsch et al., 2009), in which the gut epithelial barrier is breached in a controlled manner, leading to systemic dissemination of gut bacteria. *Ggta1*^*-/-*^ mice showed a strong survival advantage against CLP, as compared to control wild type (*Ggta1*^+*/*+^) mice (*Fig. 1A*). This was associated with a 10-10^5^-fold reduction in bacterial load, dependent upon the organ examined (*Fig. 1B*), suggesting that loss of *Ggta1* function enhances resistance to bacterial sepsis.

**Figure 1.**
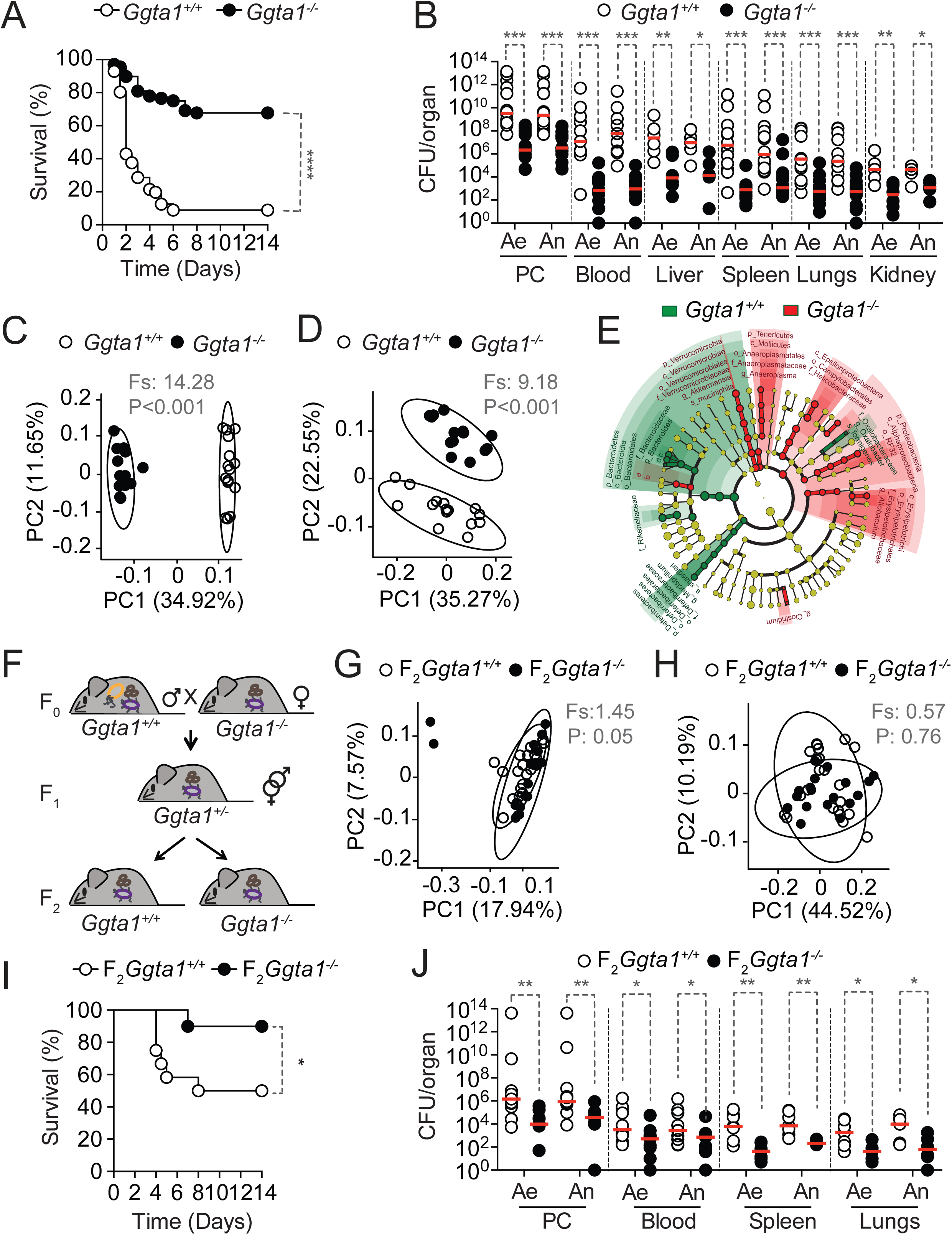
Loss of *Ggta1* function in mice enhances resistance to systemic infection by symbiotic gut bacteria. **A**) Survival after CLP of *Ggta1*^+*/*+^ (n=56) and *Ggta1*^*-/-*^ (n=68) mice; 19 experiments. **B**) Colony forming units (CFU) of aerobic (Ae) and anaerobic (An) bacteria recovered from *Ggta1*^+*/*+^ (n=5-15) and *Ggta1*^*-/-*^ (n=6-15) mice, 24 hours after CLP; 11 experiments. Principal Coordinate Analysis (PCoA) of **C**) Unweighted UniFrac distance and **D**) Weighted UniFrac distance of 16S rRNA amplicons, from fecal samples of *Ggta1*^+*/*+^ (n=15) and *Ggta1*^*-/-*^ (n=14) mice. **E**) Cladograms, from LEfSe analysis, representing taxa enriched in *Ggta1*^+*/*+^ (green) or *Ggta1*^*-/-*^ (red) mice in the same samples as (A-B). a: family_Porphyromonadaceae, b: genus_ Parabacteroides, c: species_*Bacteroides acidifaciens*, d: species_*B. ovatus*; 1 experiment. **F**) Breeding strategy for the generation of F_2_ *Ggta1*^+*/*+^ *vs. Ggta1*^*-/-*^ littermates with similar microbiota, maternally derived from *Ggta1*^*-/-*^ mice. Microbiota PCoA of **G**) Unweighted UniFrac distance and **H**) Weighted UniFrac distance of 16S rRNA amplicons, in fecal samples from F_2_ *Ggta1*^+*/*+^ (n=22) and *Ggta1*^*-/-*^ (n=18) mice generated as described in (F); 1 experiment. **I**) Survival after CLP of F_2_ *Ggta1*^+*/*+^ (n=12) and *Ggta1*^*-/-*^ (n=10) mice; 3 experiments. **J**) CFU of Ae and An bacteria of F_2_ *Ggta1*^+*/*+^ (n=5-11) and *Ggta1*^*-/-*^ (n=4-7) mice, 24 hours after CLP; 3 experiments. Symbols in (B, C, D, G, H, J) are individual mice. Red bars (B, J) show median values. P values in (A, I) calculated using log-rank test, in (C, D, G, H) using PERMANOVA test and in (B, J) using Mann-Whitney test. Peritoneal cavity (PC). *P < 0.05; **P < 0.01; ***P < 0.001; ****P < 0.0001.

The outcome of CLP is affected by the bacterial composition of the gut microbiota, which we found to be notably different between *Ggta1*^*-/-*^ and *Ggta1*^+*/*+^ mice, as determined by 16S rRNA gene sequencing analysis of the fecal microbiota for bacterial community structure (*Fig. 1C,D*), composition (*Fig. 1E*) and diversity (*Fig. S1A,B*). To dissect the contribution of host genotype *vs*. microbiota composition in conferring protection against sepsis, we used a previously established approach to normalize the microbiota between mice with different genotypes (Ubeda et al., 2012). We confirmed that vertical transmission from female *Ggta1*^*-/-*^ mice (*Fig. 1F*), normalized the bacterial composition of the gut microbiota in F_2_ littermate mice from both genotypes (*Fig. 1G-H, S1C-D*). Nevertheless, F_2_ *Ggta1*^*-/-*^ mice retained a survival advantage following CLP, when compared to F_2_ *Ggta1*^+*/*+^ mice (*Fig. 1I*). This was associated with a 10-10^2^-fold reduction in bacterial load (*Fig. 1J*), showing that loss of *Ggta1* function is sufficient *per se* to enhance resistance to bacterial sepsis. We note, however, that the relative survival advantage against CLP was higher in F_0_ *Ggta1*^*-/-*^ *vs. Ggta1*^+*/*+^ mice (*Fig. 1A,B*) harboring distinct microbiota, as compared to F_2_ *Ggta1*^*-/-*^ *vs. Ggta1*^+*/*+^ mice (*Fig. 1I,J*) harboring the same microbiota. This argues for a synergistic contribution of both the host genotype and its microbiota to enhanced protection from bacterial sepsis observed in *Ggta1*^*-/-*^ *vs. Ggta1*^+*/*+^ mice.

### Loss of *Ggta1* function in mice enhances IgG-mediated resistance to systemic polymicrobial infection

Different components of the adaptive immune systems (Kato et al., 2014), including circulating NAb that are generated in response to immunogenic bacteria in the gut microbiota (Kamada et al., 2015; Koch et al., 2016; Macpherson et al., 2018), restrain the growth of pathobionts and favor gut colonization by commensal bacteria while increasing overall microbiota diversity (Round and Palm, 2018). In keeping with this notion, the gut microbiota of *Rag2*^*-/-*^*Ggta1*^*-/-*^ mice, inheriting the microbiota from *Ggta1*^*-/-*^ mice in the absence of adaptive immunity (*Fig. S1E*), was remarkably distinct from that of *Ggta1*^*-/-*^ mice (*Fig. S1F,G*). Moreover, the microbiota of *Rag2*^*-/-*^*Ggta1*^*-/-*^ mice exhibited a marked reduction in diversity (*Fig. S1H,I*) and an enrichment of several bacteria associated with pathobiont behavior, such as Proteobacteria, *Helicobacter* and *Bacteroides* (Palm et al., 2014) (*Fig. S1J,K*). This supports the view that the adaptive immune system of *Ggta1*^*-/-*^ mice shapes the gut microbiota composition, which is consistent, albeit more pronounced, with studies in *Ggta1*^+*/*+^ mice (Barroso-Batista et al., 2015).

Circulating NAb can limit the translocation of bacterial pathobionts across the gut epithelium and consequently prevent systemic infections from triggering the onset of sepsis (Kamada et al., 2015; Zeng et al., 2016). We therefore asked whether the loss of *Ggta1* function promotes NAb-driven resistance against pathobionts present in the gut microbiota from *Rag2*^*-/-*^*Ggta1*^*-/-*^ mice. As with CLP (*Fig. 1A-B*), *Ggta1*^*-/-*^ mice were more resistant than *Ggta1*^+*/*+^ mice to intra-peritoneal (i.p.) inoculation of cecal content from *Rag2*^*-/-*^*Ggta1*^*-/-*^ mice (*Fig. 2A*), showing a 10^4^-10^7^-fold reduction in bacterial load (*Fig. 2B*). Both genotypes survived when challenged (i.p.) with a paraformaldehyde-fixed cecal inoculum (*Fig. S1L*), suggesting that loss of *Ggta1* enhances anti-bacterial resistance rather than providing protection against inflammation-driven immunopathology, a hallmark of sepsis (Rudd et al., 2020; Singer et al., 2016).

**Figure 2.**
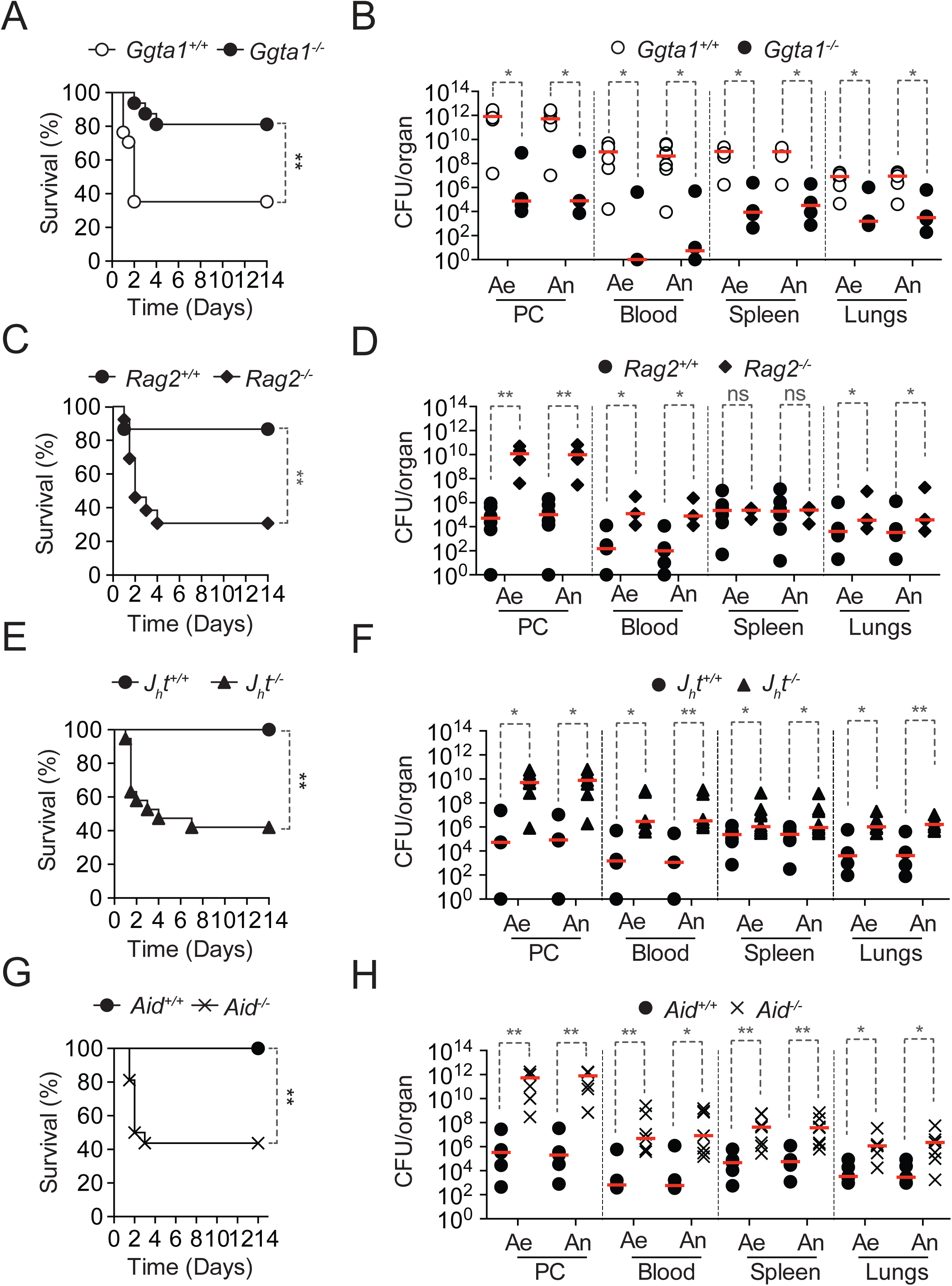
Loss of *Ggta1* function in mice enhances IgG-mediated resistance to systemic polymicrobial infection. **A**) Survival of *Ggta1*^+*/*+^ (n=16) and *Ggta1*^*-/-*^ (n=14) mice infected (*i*.*p*.) with a cecal inoculum from *Rag2*^*-/-*^*Ggta1*^*-/-*^ mice; 4 experiments. **B**) Colony forming units (CFU) of aerobic (Ae) and anaerobic (An) bacteria of *Ggta1*^+*/*+^ (n=6) and *Ggta1*^*-/-*^ (n=4) mice, 24 hours after infection; 2 experiments. **C**) Survival of *Rag2*^+*/*+^*Ggta1*^*-/-*^ (n=15) and *Rag2*^*-/-*^*Ggta1*^*-/-*^ (n=13) mice infected as in (A); 3 experiments. **D**) CFU of Ae and An bacteria of *Rag2*^+*/*+^*Ggta1*^*-/-*^ (n=5-6) and *Rag2*^*-/-*^*Ggta1*^*-/-*^ (n=4) mice, 24 hours after infection; 3 experiments. **E**) Survival of *J*_*h*_*t*^+*/*+^*Ggta1*^*-/-*^ (n=12) and *J*_*h*_*t*^*-/-*^*Ggta1*^*-/-*^ (n=19) mice infected as in (A); 4 experiments. **F**) CFU of Ae and An bacteria of *J*_*h*_*t*^+*/*+^*Ggta1*^*-/-*^ (n=4-6) and *J*_*h*_*t*^*-/-*^*Ggta1*^*-/-*^ (n=7-11) mice, 24 hours after infection; 4 experiments. **G**) Survival of *Aid*^+*/*+^*Ggta1*^*-/-*^ (n=10) and *Aid*^*-/-*^*Ggta1*^*-/-*^ (n=16) mice infected as in (A); 3 experiments. **H**) CFU of Ae and An bacteria of *Aid*^+*/*+^*Ggta1*^*-/-*^ (n=5) and *Aid*^*-/-*^*Ggta1*^*-/-*^ (n=6-7) mice, 24 hours after infection; 4 experiments. Symbols (B, D, F, H) are individual mice. Red bars (B, D, F, H) are median values. P values in (A, C, E, G) calculated with log-rank test and in (B, D, F, H) with Mann-Whitney test. Peritoneal cavity (PC). *P < 0.05; **P < 0.01; ns: not significant.

Confirming that loss of *Ggta1* enhances resistance to bacterial sepsis through mechanisms dependent upon adaptive immunity, *Rag2*^*-/-*^*Ggta1*^*-/-*^ mice were extremely susceptible to infection with the same cecal inoculum (*Fig. 2C*), carrying a 10-10^6^-fold higher bacterial load, as compared to *Ggta1*^*-/-*^ mice (*Fig. 2D*). This was also observed in *J*_*h*_*t*^*-/-*^*Ggta1*^*-/-*^ mice lacking B cells (*Fig. 2E,F*), suggesting that anti-bacterial resistance is provided by a B cell-dependent mechanism.

Although protective against bacterial infection (Ochsenbein et al., 1999), circulating IgM was not essential in this experimental system to enhance anti-bacterial resistance, as demonstrated in *µS*^*-/-*^*Ggta1*^*-/-*^ mice lacking circulating IgM (*Fig. S2A,B*). Similarly, while IgA can be critical to support mucosal immunity (Macpherson et al., 2018; Sutherland et al., 2016) and prevent the development of sepsis (Wilmore et al., 2018), it was not essential to support anti-bacterial resistance in *Iga*^*-/-*^*Ggta1*^*-/-*^ mice lacking IgA (*Fig. S2C,D*). In contrast, protection against infection with the cecal inoculum isolated from *Rag2*^*-/-*^*Ggta1*^*-/-*^ mice was lost in *Aid*^*-/-*^*Ggta1*^*-/-*^ mice lacking immunoglobulin (Ig) class-switch recombination, somatic hypermutation and affinity maturation (Muramatsu et al., 2000) (*Fig. 2G*). This was associated with a 10^3^-10^6^-fold increase in bacterial load, as compared to *Ggta1*^*-/-*^ mice (*Fig. 2H*), suggesting that the survival advantage against bacterial sepsis in *Ggta1*^*-/-*^ mice relies on the presence of circulating NAb that must undergo class-switch recombination towards the IgG isotype.

Immunoglobulin class-switch, somatic hypermutation and affinity maturation are largely T cell-dependent, suggesting that T cells might support resistance to infection in *Ggta1*^*-/-*^ mice. However, we observed that *Tcrβ*^*-/-*^*Ggta1*^*-/-*^ (*Fig. S2E,F*) and *Tcrδ*^*-/-*^ *Ggta1*^*-/-*^ (*Fig. S2G,H*) mice, lacking α/β and γ/δ T cells respectively, were resistant to sepsis, similarly to *Ggta1*^*-/-*^ mice. This suggests that the protective circulating IgG NAb in *Ggta1*^*-/-*^ mice are generated in a T-cell independent manner.

### The protective effect of IgG NAb acts irrespectively of αGal recognition

We then asked whether circulating IgG NAb from *Ggta1*^*-/-*^ mice are sufficient *per se* to confer resistance to infection. Upon adoptive transfer at the same dosage to *Rag2*^*- /-*^*Ggta1*^*-/-*^ mice, only circulating IgG NAb purified from *Ggta1*^*-/-*^ mice, but not from *Ggta1*^+*/*+^ mice, were protective against infection (*Fig. 3A*). Since the concentration of circulating IgG in naive *Ggta1*^*-/-*^ mice that are protected from infection, was similar to that of *Ggta1*^+*/*+^ mice, that are not protected (*Fig. S3A*), this suggests that IgG Nab from *Ggta1*^*-/-*^ mice have an enhanced capacity to confer protection against bacterial infection, as compared to the IgG NAb from *Ggta1*^+*/*+^ mice.

**Figure 3.**
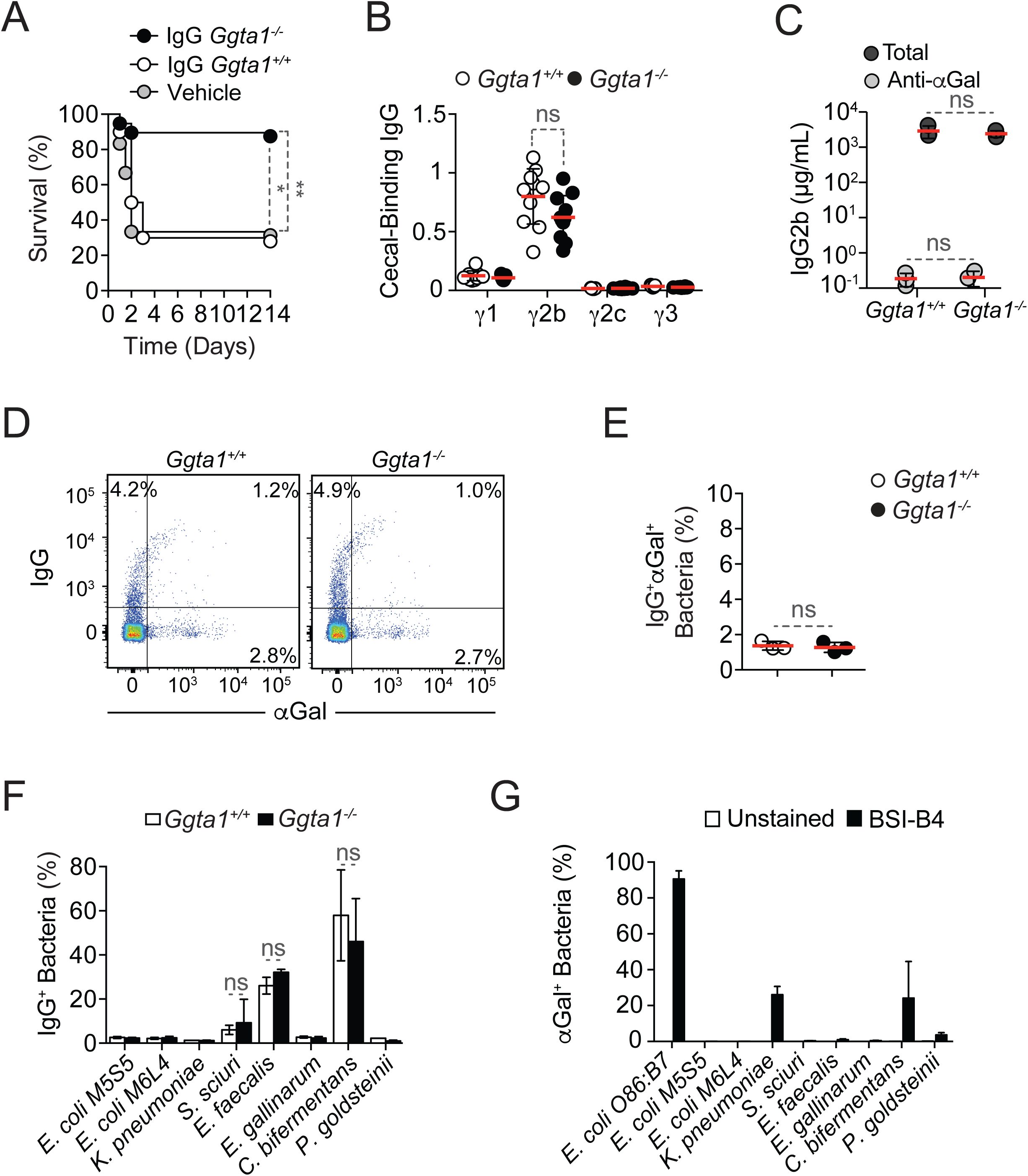
The protective effect of IgG NAb acts irrespectively of αGal recognition. **A**) Survival of *Rag2*^*-/-*^*Ggta1*^*-/-*^ mice after transfer of IgG purified from *Ggta1*^*-/-*^ mice (n=15), *Ggta1*^+*/*+^ mice (n=10) or vehicle (PBS; n=6), 24 hours before infection (*i*.*p*.) with a cecal inoculum from *Rag2*^*-/-*^*Ggta1*^*-/-*^ mice; 4 experiments. **B**) Relative absorbance of IgG sub-classes in the serum of *Ggta1*^+*/*+^ (n=11) and *Ggta1*^*-/-*^ (n=11) mice, binding to cecal extract from *Rag2*^*-/-*^*Ggta1*^*-/-*^ mice, determined by ELISA; representative of 3 experiments. **C**) Concentration of total IgG2b and anti-αGal IgG2b in IgG from *Ggta1*^+*/*+^ or *Ggta1*^*-/-*^ mice (n=3). **D**) Representative plots showing *Rag2*^*-/-*^ *Ggta1*^*-/-*^ cecal bacteria co-stained with IgG from *Ggta1*^+*/*+^ or *Ggta1*^*-/-*^ mice and BSI-B4 lectin (αGal). **E**) Quantification of IgG^+^αGal^+^ bacteria in the same samples as in (C) (n=3); 3 experiments. **F)** Percentage of *in vitro*-grown bacteria isolated from the mouse microbiota bound by IgG purified *Ggta1*^+*/*+^ and *Ggta1*^*-/-*^ mice. Data shown was pooled from N=3-4 independent experiments. **G**) Quantification of αGal expression by *in vitro*-grown species of bacteria as indicated; 3 independent experiments. Symbols in (B) are individual mice and in (C, E) are independent IgG preparations. Red lines in (B, C, E) are mean values. Error bars in (B, C, E, F, G) correspond to SD. P values in (A) calculated using log-rank test, in (B, E) using Mann-Whitney test and in (C, F) by 2-way ANOVA using Sidak’s multiple comparison test. *P < 0.05; **P < 0.01; ns: not significant.

We reasoned that the enhanced capacity of IgG NAb from *Ggta1*^*-/-*^ mice to confer protection against bacterial infection might be due to changes in IgG subclass composition, enhancing overall effector function (Lu et al., 2018). Alternatively, differences in the repertoire of circulating IgG NAb could enable the recognition of different bacterial antigens including, αGal-like glycans.

We found that the bacterium-binding IgG NAb from *Ggta1*^*-/-*^ and *Ggta1*^+*/*+^ mice were almost exclusively from the IgG2b subclass (*Fig. 3B*), analogous to previous reports in *Ggta1*^+*/*+^ mice (Zeng et al., 2016). This excludes the possibility that increased resistance to bacterial infection provided by IgG NAb from *Ggta1*^*-/-*^ *vs. Ggta1*^+*/*+^ mice is due to differences in relative IgG subclass composition.

A number of bacteria in the gut microbiota express αGal-like glycans (Montassier et al., 2019), suggesting that αGal-specific NAb might be protective against systemic infection by these bacteria. When maintained under specific pathogen-free conditions however, circulating αGal-specific NAb accounted for less than 0.005% of circulating IgG2b NAb from *Ggta1*^*-/-*^ and *Ggta1*^+*/*+^ mice (*Fig. 3C*), which is consistent with previous reports (Galili et al., 1984; Yilmaz et al., 2014). In keeping with this observation, circulating IgG2b NAb from both *Ggta1*^*-/-*^ and *Ggta1*^+*/*+^ mice recognized only a negligible, but similar proportion of αGal-expressing bacteria in the infectious inoculum (*Fig. 3D,E, S3B*). Taken together, this suggests that loss of *Ggta1* enhances protection against bacterial sepsis via a mechanism that acts irrespectively of αGal-specific immunity.

We then asked whether circulating IgG NAb from *Ggta1*^*-/-*^ and *Ggta1*^+*/*+^ mice recognized to the same or different extent individual bacterial strains isolated from mouse microbiota. We found that circulating IgG NAb from *Ggta1*^*-/-*^ and *Ggta1*^+*/*+^ mice recognized these bacteria to the same extent (*Fig. 3F, S3C*), irrespectively of αGal expression by the targeted bacteria (*Fig*.*3G, S3C,D*). This is illustrated for *E. faecalis*, which does not express αGal, and *C. bifermentans* that expresses relatively low levels of αGal, when compared to *E. coli* O86:B7 that expresses high levels of αGal (*Fig*.*3G, S3C,D*). This suggests that the enhanced protective effect exerted by the circulating IgG NAb from *Ggta1*^*-/-*^ *vs. Ggta1*^+*/*+^ mice does not rely on the recognition of αGal-like glycans in the targeted bacteria.

### Loss of *Ggta1* function does not affect bacterial recognition by IgG NAb

Having established that protection against bacterial sepsis was independent of αGal-specific immunity, we asked whether circulating IgG NAb from *Ggta1*^*-/-*^ mice recognized distinct bacteria, when compared to IgG NAb from *Ggta1*^+*/*+^ mice. The percentage of bacteria recognized in the cecal inoculum used for infection by IgG NAb from *Ggta1*^*-/-*^ *vs. Ggta1*^+*/*+^ mice was indistinguishable, as assessed *in vitro* by staining of bacteria (*Fig. 4A*) and *in vivo* by detecting IgG-bound bacteria recovered from the peritoneal cavity after infection with this inoculum (*Fig. 4B*). Relative amount of IgG bound *per* bacterium was also similar in both experimental settings (*Fig. S4A,B*). Co-staining of the same infectious inoculum with IgG NAb from *Ggta1*^*-/-*^ and *Ggta1*^+*/*+^ mice, conjugated to different fluorophores, demonstrated that these NAb recognized the same bacteria (*i*.*e*. >97% similar), as illustrated by flow cytometry (*Fig. 4C,D*). Moreover, the pattern of bacterial recognition obtained using IgG NAb from the same genotype, but conjugated to different fluorophores, was indistinguishable from that obtained using IgG NAb from different genotypes (*S4C-F*). Recognition of largely overlapping bacteria by IgG NAb from *Ggta1*^*-/-*^ *vs. Ggta1*^+*/*+^ mice was confirmed by 16S rRNA analysis of IgG-bound (*Fig. 4E-F, S4G*) and non-IgG-bound bacteria (*Fig. 4G-H, S4H*). These observations suggest that circulating IgG NAb from both genotypes recognize the same bacteria in the infectious inoculum, and do so to the same extent. Therefore, the enhanced resistance to bacterial sepsis provided by the IgG NAb from *Ggta1*^*-/-*^ *vs. Ggta1*^+*/*+^ mice is likely not due to differences in the bacteria recognized.

**Figure 4.**
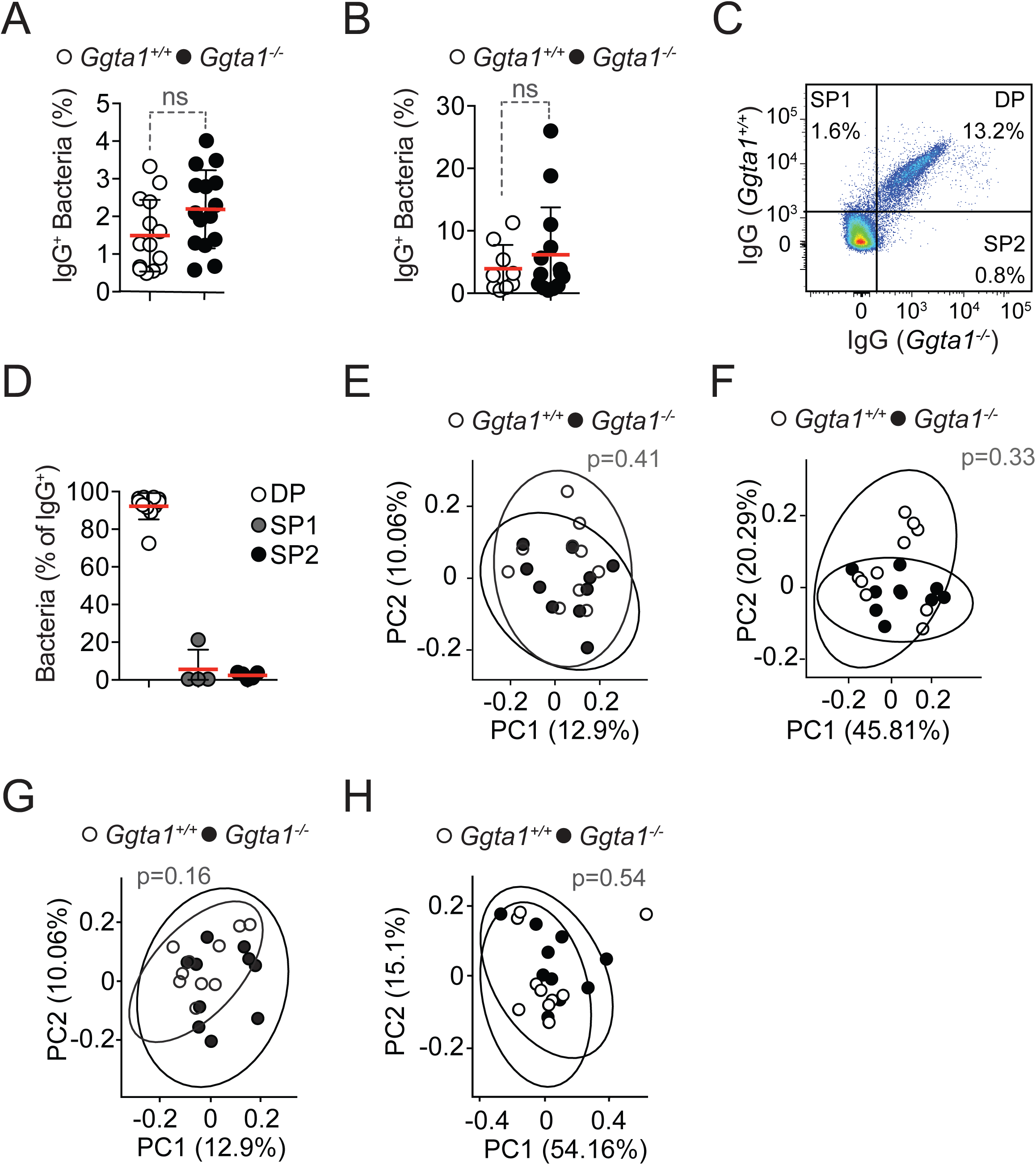
Loss of *Ggta1* function does not alter bacterial recognition by IgG NAb. **A**) Percentage of IgG^+^ bacteria in the *Rag2*^*-/-*^*Ggta1*^*-/-*^ cecal content, after incubation with serum from *Ggta1*^+*/*+^ (n=14) and *Ggta1*^*-/-*^ (n=15) mice; 3 experiments. **B**) Percentage of IgG^+^ bacteria in the peritoneal lavage of *Ggta1*^+*/*+^ (n=9) and *Ggta1*^*-/-*^ (n=14) mice, 3 hours after infection (*i*.*p*.) with a cecal inoculum from *Rag2*^*-/-*^*Ggta1*^*-/-*^ mice; 3 experiments. **C**) Representative plot showing *Rag2*^*-/-*^*Ggta1*^*-/-*^ cecal bacteria co-stained with IgG from *Ggta1*^+*/*+^ and *Ggta1*^*-/-*^ mice. **D**) Percentage of double positive (DP) *vs*. single positive (SP) bacteria among all IgG^+^ bacteria in the same samples as in (C) (n=11); 6 experiments. **E-H**) Principal Coordinates Analysis of **E**) Unweighted UniFrac and **F**) Weighted UniFrac distance of IgG^+^ bacteria and **G**) Unweighted UniFrac and **H**) Weighted UniFrac distance of IgG^-^ bacteria in the peritoneal lavage of *Ggta1*^+*/*+^ (n=10) and *Ggta1*^*-/-*^ (n=10) mice, 3 hours after infection with a cecal inoculum from *Rag2*^*-/-*^*Ggta1*^*-/-*^ mice; 2 experiments. Symbols in (A, B, E, F, G, H) are individual mice and in (D) are independent *Rag2*^*-/-*^*Ggta1*^*-/-*^ cecal preparations. Red lines (A, B, D) are mean values. Error bars in (A, B, D) correspond to SD. P values in (A, B) calculated using Mann-Whitney test and in (E-H) using PERMANOVA test. ns: not significant.

### Loss of αGal expression enhances IgG effector function

We then reasoned that the enhanced protective effect exerted by the IgG NAb from *Ggta1*^*-/-*^ *vs. Ggta1*^+*/*+^ mice might be due to a corresponding enhancement of IgG effector function. To test this hypothesis, we used the bacterial strain, *E. faecalis* that lacks αGal but is recognized to a similar extent by IgG NAb from *Ggta1*^*-/-*^ *vs. Ggta1*^+*/*+^ mice (*Fig. 3F,G, S3C,D*). Naïve *J*_*h*_*t*^*-/-*^*Ggta1*^*-/-*^ mice, lacking circulating Ig, were challenged (i.p.) with *E. faecalis*, opsonized, or not, by circulating IgG NAb purified from *Ggta1*^*-/-*^ or *Ggta1*^+*/*+^ mice. Recruitment of neutrophils (CD11b^+^Ly6G^high^) into the peritoneal cavity was enhanced only when *E. faecalis* was opsonized with the IgG from *Ggta1*^*-/-*^, but not from *Ggta1*^+*/*+^ mice, as compared to naïve *J*_*h*_*t*^*-/-*^*Ggta1*^*-/-*^ mice (*Fig. 5A, S5A*). This was associated with an increase in the number of peritoneal neutrophils containing *E. faecalis* (*Fig. 5B, S5B,C*), probably due to increased, albeit borderline significant, IgG-dependent phagocytosis (*Fig. 5C, S5C*). This suggests that upon bacterial recognition, the relative capacity of IgG NAb from *Ggta1*^*-/-*^ mice to promote bacterial phagocytosis by neutrophils is enhanced, when compared to IgG from *Ggta1*^+*/*+^ mice. Moreover, this effect acts independently of αGal-recognition in the targeted bacteria, suggesting that the loss of *GGTA1* function enhances the effector function of circulating IgG NAb, irrespectively of the epitope recognized.

**Figure 5.**
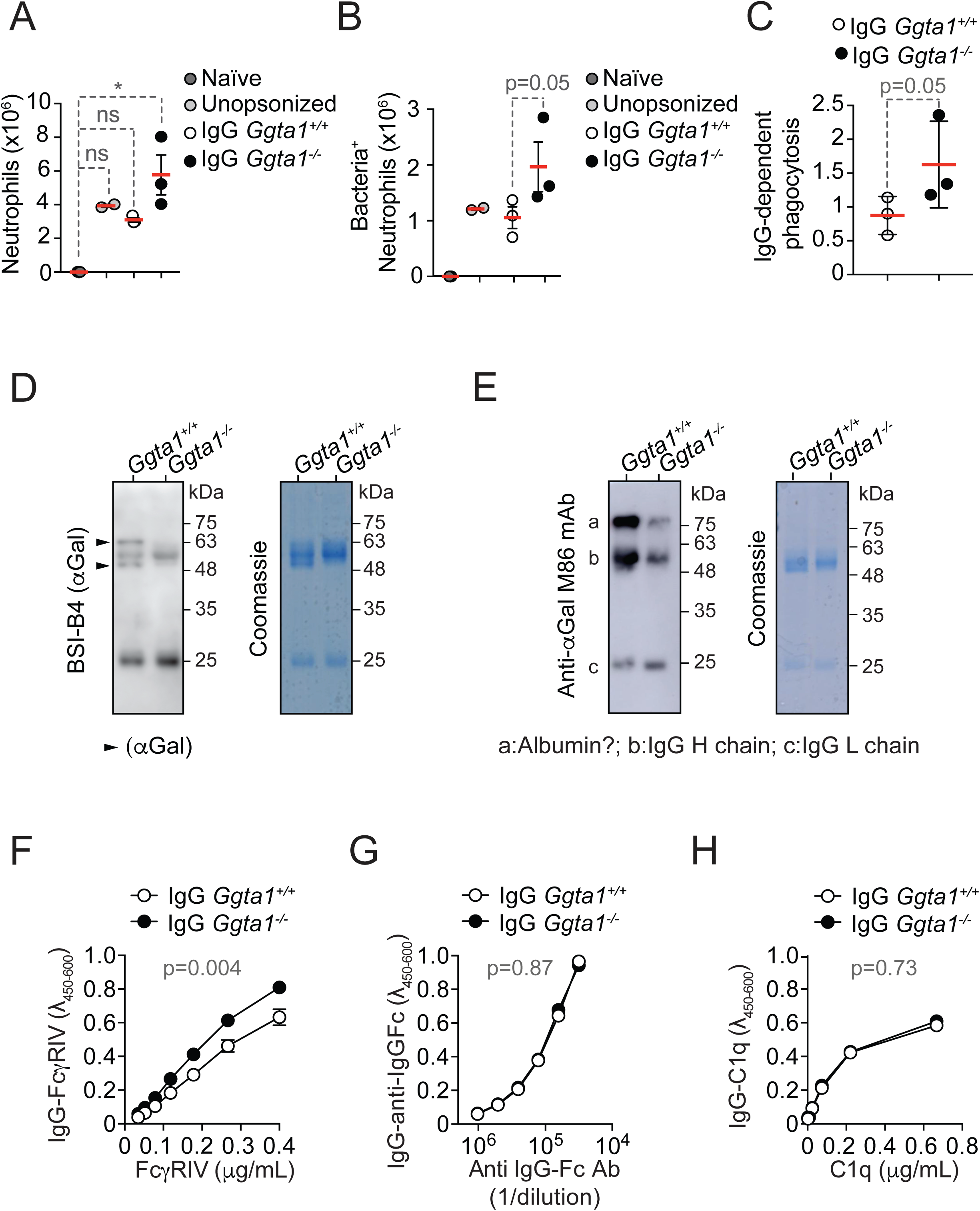
Loss of αGal expression enhances IgG effector function. **A**) Total number of infiltrating CD11b^+^Ly6G^high^ neutrophils recovered from the peritoneal cavity of *J*_*h*_*t*^*-/-*^*Ggta1*^*-/-*^ mice, either naive (dark gray circles; n=3) or 3 hours after injection of *E. faecalis*, unopsonized (light gray circles; n=2), or opsonized with IgG from *Ggta1*^+*/*+^ (white circles; n=3) or *Ggta1*^-/-^ (black circles; n=3) mice; 1 experiment. **B)** Total number of bacteria-containing neutrophils in the same samples as in (A). **C**) IgG-dependent phagocytosis index calculated as a ratio of bacteria-containing neutrophils in IgG-opsonized groups over the unopsonized group, in the same samples as in (A). **D-E)** Detection of αGal in IgG from *Ggta1*^+*/*+^ and *Ggta1*^*-/-*^ mice, using **D)** BSI-B4 lectin and **E**) Anti-αGal M86 mAb. SDS gel is shown as loading control. Representative of 6 independent IgG preparations. **F**) Relative binding to FcγRIV and **G**) Anti-IgG by *Ggta1*^+*/*+^ (n=6) and *Ggta1*^*-/-*^ (n=6) purified IgG. **H**) Relative binding to mouse C1q by *Ggta1*^+*/*+^ (n=7) and *Ggta1*^*-/-*^ (n=3) purified IgG, where n corresponds to independent IgG preparations. Data is representative of 1-3 independent experiments. Symbols in (A, B, C) are individual mice. Red lines (A, B, C) are mean values. Error bars in (A, B, C) correspond to SD and in (F, G, H) to SEM. P values in (A) calculated using a one-tailed Kruskal-Wallis test with Dunn’s correction for multiple comparisons, in (B, C) using Mann-Whitney test and in (F-H) using 2-Way ANOVA with Sidak’s multiple comparison test. *P < 0.05; ns: not significant.

It is well established that the relative composition of the glycan structures, N-linked to Asn297 of the constant heavy (H) chain of the IgG Fc domain, can have a considerable impact upon IgG-FcγR binding and downstream IgG effector functions (Anthony et al., 2012; Dekkers et al., 2017; Wang and Ravetch, 2019). Having confirmed the presence of αGal in the H chain of circulating IgG NAb from *Ggta1*^+*/*+^ mice (de Haan et al., 2017), but not from *Ggta1*^*-/-*^ mice (*Fig. 5D,E, Fig. S5D*), we asked whether αGal modulates IgG effector function in a manner that impacts on resistance to bacterial sepsis.

We found that the relative binding of mouse FcγRIV to IgG NAb from *Ggta1*^*-/-*^ mice was enhanced, as compared to IgG NAb from *Ggta1*^+*/*+^ mice (*Fig. 5F*). In contrast, there were no differences in the relative binding of an anti-IgG Fc Ab to IgG from either *Ggta1*^*-/-*^ or *Ggta1*^+*/*+^ mice in the same assay (*Fig. 5G*). This shows that increased binding of mouse FcγRIV to IgG from *Ggta1*^*-/-*^ or *Ggta1*^+*/*+^ mice in this assay is specific. Further strengthening this notion, the relative binding of other mouse FcγR to IgG NAb from *Ggta1*^*-/-*^ *vs. Ggta1*^+*/*+^ mice was indistinguishable, as illustrated for FcγRI (*Fig. S5E*), FcγRIIb (*Fig. S5F*), FcγRIII (*Fig. S5G*) and FcRn (*Fig. S5H*). There was also no difference in the relative binding of mouse complement component 1q (C1q) to IgG from *Ggta1*^*-/-*^ *vs. Ggta1*^+*/*+^ mice (*Fig. 5H*). Overall, this suggests that when present in the glycan structure associated to the IgG H chain, αGal hinders IgG Fc recognition by FcγRIV. This is consistent with bacteria-reactive IgG NAb being exclusively IgG2b (*Fig. 3B*), an IgG sub-class recognized preferentially by FcγRIV (Nimmerjahn et al., 2010), the mouse orthologue of human FcγRIIIA (Nimmerjahn et al., 2005). We thus infer that the absence of αGal from the Fc-glycan structure of IgG enhances the effector function of circulating IgG NAb and presumably therefore, their protective effect against bacterial sepsis.

### Loss of *Ggta1* function precipitates reproductive senescence

Loss of *GGTA1* function was a sporadic event in mammalian evolution, almost unique to Old World primates (Galili et al., 1988b; Galili and Swanson, 1991). This suggests that in most other mammals, even when considering its associated survival advantage against infection, the loss of *GGTA1* is associated with a major fitness cost, *i*.*e*. an evolutionary trade-off (Stearns and Medzhitov, 2015). Consistent with this notion, we observed a marked reduction in the total number of offspring produced by *Ggta1*^*-/-*^ compared to *Ggta1*^+*/*+^ mice throughout their reproductive lifespan (*Fig. 6A*). While consistent with a previous report suggesting that αGal partakes in mammalian reproduction (Bleil and Wassarman, 1988), other studies do not report a conclusive, physiological role for αGal in this process (Thall et al., 1995).

**Figure 6.**
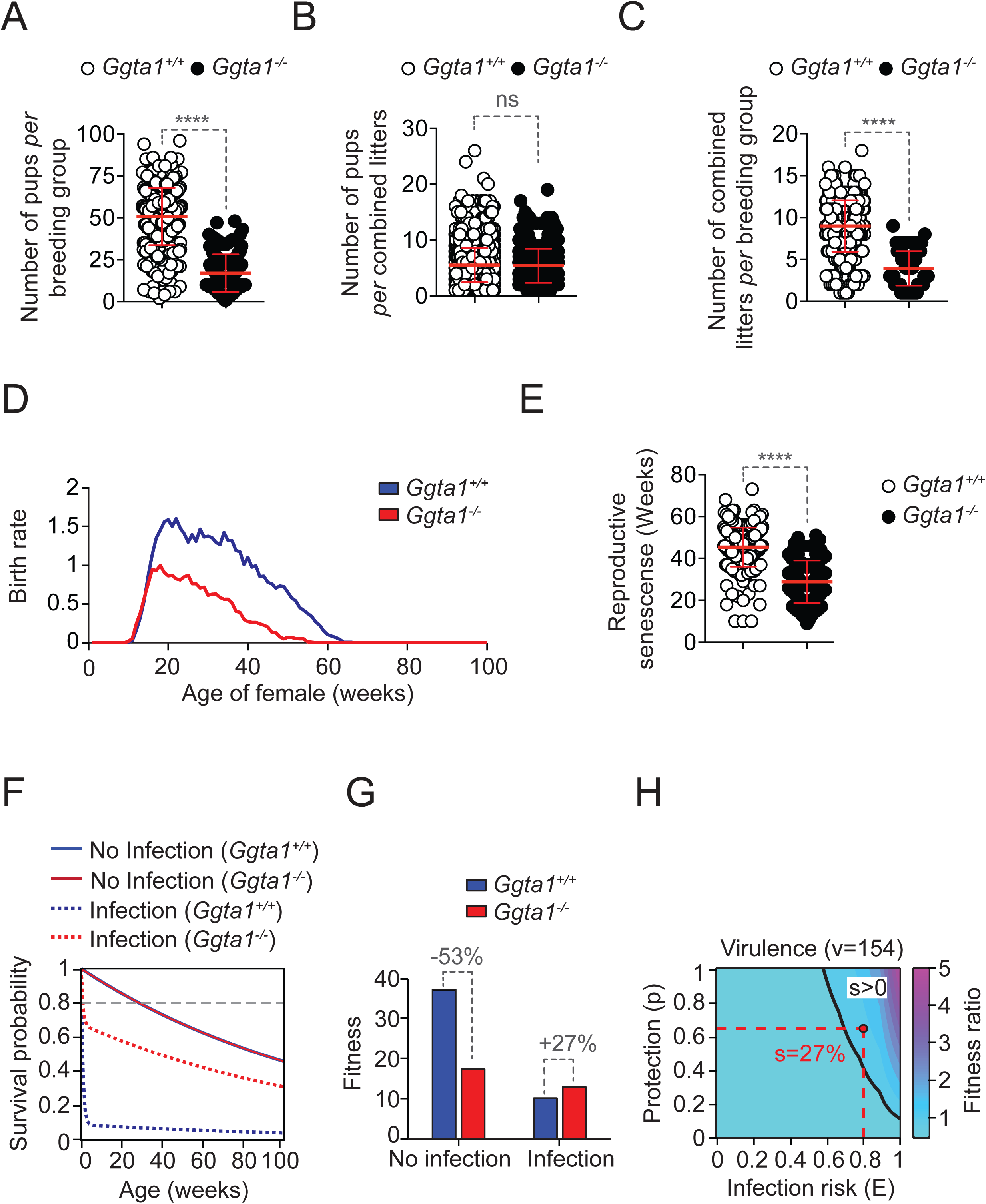
The increase in fitness provided by the survival advantage against infection can outweigh the reproductive fitness cost associated with loss of *Ggta1*. **A**) Cumulative number of offspring (pups) produced over the reproductive lifespan of *Ggta1*^+*/*+^ (n=432) and *Ggta1*^*-/-*^ (n=135) trio breeding groups, over a period of five years. **B**) Number of offspring at the time of weaning *per* combined litter from the same trio breeding groups as in (A). **C**) Number of combined litters produced over the reproductive lifespan of the same trio breeding groups as in (A). **D**) Birth rate as a function of age of females in the same trio breeding groups as in (A). **E**) Age of females at the time of the last living combined litter. **F**) Survival probability of non-infected *vs*. infected *Ggta1*^*-/-*^ and *Ggta1*^+*/*+^ mice. Dashed gray line depicts a scenario assuming high and constant exposure to infection (i.e. 80% probability at any age). **G**) Overall fitness of *Ggta1*^*-/-*^ and *Ggta1*^+*/*+^ mice, under conditions of no exposure or high exposure to infection. The reproductive cost of *Ggta1* deletion and survival advantage upon infection calculated in (F) is multiplied with the birth rate functions calculated in (D). **H**) Contour plot in which lifetime exposure to infection (*E*), assumed as constant over age, is varied along the x-axis and the magnitude of protection in *Ggta1*^*-/-*^ relative to *Ggta1*^+*/*+^ (*p*) is varied along the y-axis. The contour plot maps combinations of *E* and *p* to theoretical model-predictions for the fitness ratio (>1 favoring *Ggta1*^*-/-*^, and <1 favoring *Ggta1*^+*/*+^). Black line indicating fitness ratio=1 sets the threshold for positive selective advantage (*s>0*) of *Ggta1*^*-/-*^ *vs. Ggta1*^+*/*+^ genotype. Red lines depict a scenario in which loss of *Ggta1*, conferring protection *p* of 64%, against highly virulent pathogens (*v= 154*) reaches a selective advantage *(s*) of 27%, despite the fitness cost shown in (D), when exposure is high (*E=80%*). Error bars in (A, B, C and E) correspond to mean ± SD. P values in (A, B, C and E) calculated with Mann-Whitney test. ****P < 0.0001, ns: not significant.

We found that *Ggta1*^*-/-*^ mice produced normal numbers of viable offspring *per* litter (*Fig. 6B*), which is in keeping with previously published data (Thall et al., 1995). However, we also found that *Ggta1*^*-/-*^ mice had a reduced cumulative reproductive output compared to *Ggta1*^+*/*+^ mice due to a reduction in the total number of litters produced during their reproductive lifespan (*Fig. 6C*). This phenotype had not previously been reported (Thall et al., 1995).

The birth rate as a function of age in *Ggta1*^*-/-*^ females was reduced compared to that of *Ggta1*^+*/*+^ females (*Fig. 6D*). Age-specific birth rate is used herein to define the product of: probability of birth at any age, mean number of pups produced *per* combined litter at any age, and number of combined litters produced *per* breeding group during its reproductive lifespan. Average age at which *Ggta1*^*-/-*^ females produced their last viable litter was lower, when compared to *Ggta1*^+*/*+^ females (*Fig. 6E*), suggesting that loss of *Ggta1* function precipitates the onset of reproductive senescence.

### Protection against infection can outweigh the reproductive fitness cost associated with loss of *Ggta1*

We then estimated the likelihood of *GGTA1* loss-of-function mutations being naturally selected and fixed in populations when taking into account their associated reproductive fitness cost. We developed a mathematical model (*See Methods*) that translates host survival advantage against infection and reproductive cost into a combined measure of fitness (*i*.*e*. lifetime reproductive success). This mathematical model takes into account the natural mortality rate of non-infected mice under laboratory conditions (Kunstyr and Leuenberger, 1975), and the empirically-observed survival advantage against infection associated with *Ggta1* deletion (*Fig. 1A*). This model predicts a marked increase in the probability of survival of *Ggta1*^*-/-*^ compared to *Ggta1*^+*/*+^ mice, under conditions of constant high exposure to infection (i.e., ≥80%), if such an advantage is present at any given age (*Fig. 6F*).

To test whether such a survival advantage could outweigh the observed trade-off in reproductive output, we incorporated the reproductive fitness cost associated with *Ggta1* deletion into this model (*See Methods*). Computing host lifetime reproductive success as a combined integral of age-specific birth rate, weighted by age-specific survival, we derived a global fitness measure that could be compared across the two host genotypes. Under defined biological scenarios, the positive fitness effect calculated from protection from infection over the lifetime outweighed the reproductive fitness cost, leading to a higher overall fitness (*Fig. 6G*). This is revealed by a positive selection coefficient (*s>0*) in some regions of the parameter space (*Fig. 6H*), indicating superior fitness of hosts carrying *Ggta1* deletion, relative to wild type. This was only achieved under conditions of high exposure to virulent pathogens against which the loss of *Ggta1* provides robust protection (*Fig. 6H, Fig. S6A-D*).

## DISCUSSION

Humans carry a *GGTA1* pseudogene, carrying two frame-shift mutations and an exon deletion, which probably occurred in ancestral apes and Old World primates about 28 million years ago (Galili et al., 1987; Galili and Swanson, 1991). We propose that these loss-of-function mutations were naturally selected and fixed in different populations independently, owed to a robust associated fitness advantage against the development of sepsis, an often-lethal outcome of infection by different types of virulent pathogens that remains a major global cause of human morbidity and mortality (Rudd et al., 2020; Singer et al., 2016).

Our findings support the notion that the fitness advantage against infection, provided by the loss of *GGTA1* function, emerged from the “removal” of αGal from IgG-associated glycan structures, increasing IgG effector function and improving resistance to bacterial sepsis. This notion is supported by several independent observations in *Ggta1*-deficient mice, which similar to humans, fail to express αGal. First, *Ggta1*-deficient mice are more resistant to systemic bacterial infections originating from the gut microbiota and leading to sepsis, when compared to wild type controls (*Fig. 1,2*). Second, the survival advantage of *Ggta1*-deficient mice against bacterial sepsis relies on circulating IgG NAb recognizing bacteria in the gut microbiota (*Fig. 2,3*). Third, circulating IgG NAb confer protection against bacterial sepsis (*Fig. 3A*) irrespectively of αGal recognition (*Fig. 3D-G*). Fourth, bacterial recognition by circulating IgG NAb from *Ggta1*-deficient mice is indistinguishable from that of IgG NAb from wild type mice (*Fig. 4*). Fifth, lack of αGal in the glycan structure of IgG NAb from *Ggta1*-deficient mice increases IgG effector function, via a mechanism associated with enhanced binding to FcγRIV (*Fig. 5*), the mouse orthologue of the human FcγRIIIA (Nimmerjahn et al., 2005), which plays a critical role in driving IgG effector function (Shields et al., 2002).

Under the experimental conditions used in our study, the overwhelming majority of the circulating IgG NAb recognizing bacteria in the infectious cecal inoculum were from the IgG2b subclass (*Fig. 3B*). FcγRIV is the main FcγR recognizing IgG2b (Nimmerjahn et al., 2005), probably explaining why IgG NAb from *Ggta1*-deficient mice increase their binding specifically towards FcγRIV, but not to other FcγR (*Fig. 5*). This does not preclude however, αGal from modulating the binding of different IgG subclasses to their corresponding FcγR (Anthony et al., 2012; Dekkers et al., 2017; Wang and Ravetch, 2019).

Our findings question to some extent the previous accepted notion that the evolutionary advantage conferred by the loss of *GGTA1* function was driven essentially by the emergence of protective immunity against αGal-expressing pathogens (Soares and Yilmaz, 2016; Takeuchi et al., 1996; Yilmaz et al., 2014). While this trait was likely naturally selected on the basis of improved resistance against a number of pathogens that express αGal provided by circulating IgM NAb (Soares and Yilmaz, 2016; Takeuchi et al., 1996; Yilmaz et al., 2014), the selective advantage provided by enhancing IgG NAb effector function probably improved resistance to a larger spectrum of pathogens, irrespectively of αGal recognition. Presumably, this is equally, if not more, important to explain the evolutionary advantage conferred by the loss of *GGTA1* function.

Despite its significant benefit against infection, the loss of *GGTA1* function is a sporadic event in mammalian evolution, almost exclusive to Old World primates (Galili et al., 1988b; Galili and Swanson, 1991). This suggests that the fitness advantage against infection is linked to a decrease in fitness through a correlated trait (Stearns, 1989). At the population level, genetic trade-offs are explained by negative coupling of traits over life history, such that one trait increases fitness early in life, while another trait is detrimental later on (Williams, 1957). We propose that the trade-off associated with loss of *GGTA1* function is the emergence of reproductive senescence late in life (*Fig. 6*). Of note, reproductive senescence is a distinguishing feature of Old World primates, in which *GGTA1* function is impaired, compared to New World primates, in which *GGTA1* is functional (Hearn, 1983).

We have estimated the likelihood of *GGTA1* loss-of-function mutations being naturally selected and fixed in populations on the basis of their associated fitness advantage against infection *vs*. reproductive fitness cost. A mathematical model that computes lifetime reproductive success based on host survival advantage against infection and reproductive cost, suggests that the selective pressure imposed by sepsis can fix *GGTA1* loss-of-function when both exposure to and virulence of the pathogen are high (*Fig. 6*). This supports the hypothesis that protection from infection by highly virulent pathogens associated with the development of sepsis might have led to a “catastrophic-selection” of ancestral primates, whereby mutant offspring lacking a functional *GGTA1* survived and replaced the parental populations (Galili, 2019).

Although the mathematical model used in this study was parameterized by empirical data obtained from mice maintained under laboratory conditions, its biological structure speaks to general scenarios, where trade-offs between protection from infection and reproduction are at play. With such a quantitative framework of lifetime reproductive success, alternative combinations of functions and parameters can be integrated to explore selection of traits that simultaneously affect reproduction and survival. This includes protective immunity against virulent pathogens expressing αGal (Soares and Yilmaz, 2016) or possibly female immunity against paternal αGal, as possible contributors to the rapid fixation of *GGTA1* loss-of-function mutations similar to other human specific loss-of-function mutations, such as in the CMP-*N*-acetylneuraminic acid hydroxylase (*CMAH*) gene (Ghaderi et al., 2011).

An evolutionary implication of our findings is that Old World primates, including humans, appear to be at a higher risk of developing sepsis in response to bacterial infection. In strong support of this notion, humans are more susceptible to develop septic shock when challenged by bacterial lipopolysaccharide (LPS), as compared to other mammalian lineages (Chen et al., 2019). Whether this can be explained by intrinsic characteristics of human immunity and/or the capacity to establish disease tolerance to infection (Martins et al., 2019; Medzhitov et al., 2012), is unclear. This is consistent however with the idea that, similar to *GGTA1*, humans carry other loss-of-function mutations associated with enhanced immune resistance to bacterial infection (Olson, 1999; Wang et al., 2006). Considering that genetic programs driving resistance and disease tolerance to infection are negatively correlated (Raberg et al., 2007), mutations increasing resistance might carry, as a trade-off, a reduced capacity to establish disease tolerance to infection.

In conclusion, our findings support the idea that the positive selection of *GGTA1* loss-of*-*function mutations in the common ancestor of Old World primates was propelled by an overall enhancement in IgG effector function, providing resistance against infection by gut bacteria pathobionts that would otherwise lead to the development of sepsis. This provided a survival advantage against infection by a broad range of pathogens, likely outweighing the trade-off imposed by the emergence of reproductive senescence and lower reproductive output, potentially explaining why loss of *GGTA1* was a rare event, which occurred almost exclusively in Old World primates, including humans.

## ACKNOWLEDGEMENTS

We thank L. Barreiro (U. Chicago), J. Howard, I. Gordo, K. Xavier, J. Demengeot, L. Chikhi, S. Ramos (Instituto Gulbenkian de Ciência) for critical review of the manuscript, M. Zeng (Weill Cornell Medicine) for providing mouse microbiota bacterial strains, M.E. Conner (Baylor College of Medicine) for the *Iga*-deficient mice, R. Knight, G. Humphrey and L. Marotz (University of California, San Diego) for help in bacterial sorting experiments, T. Paixão for data analysis and constructive reviewing of the mathematical model, K. Xavier and A.R. Oliveira for help in 16S sequencing, F. Braza, M. Monteiro and V. Martins for Flow Cytometry analyses, P.B. Amador for help with GF mice and Ab analyses, A. Regalado for IgG purification and conjugation, A. Vieira, J. Bom and A. Ribeiro (all IGC) for reproductive output data. S. S. is supported by Fundação para a Ciência e Tecnologia (FCT; SFRH/BD/52177/2013), J.A.T. by an ESCMID Research Grant and FCT (SFRH/BPD/112135/2015), S.W. by DFG Excellence Cluster EXS 2051 and CSCC, Jena University Hospital (BMBF 01EO1502), E.G by FLAD/NSF 274/2016 and M.P.S. by the Gulbenkian, “La Caixa” (HR18-00502) and Bill & Melinda Gates (OPP1148170) Foundations and FCT (5723/2014 and FEDER029411).

## AUTHOR CONTRIBUTIONS

S.S. contributed critically to study design, and performed most experimental work and data analyses. J.A.T. contributed to study design, performed experimental work and data analyses. S.W. made the original observation that loss of *Ggta1* increases resistance to sepsis with B.Y. D.S. and M.T. performed 16S rRNA sequencing analysis. B.Y., S.R. and S.C. generated mouse strains, maintained breeding colonies and gathered reproductive output data with S.S. E.G. performed mathematical modeling for integrated fitness over lifespan. G.N. provided mouse microbiota bacterial strains and contributed intellectually. M.P.S. drove the study design and wrote the manuscript with S.S., with contributions from J.A.T.

## DECLARATION OF INTERESTS

The authors declare no competing interests.

## STAR METHODS

Detailed methods are provided in the online version of this paper and include the following:

- **KEY RESOURCES TABLE**
- **CONTACT FOR REAGENTS AND RESOURCE SHARING**
- **EXPERIMENTAL MODEL AND SUBJECT DETAILS**
  - Mice
  - Breeding experiments
  - Reproductive output
  - Cecal Ligation and Puncture
  - Cecal Slurry Injection

- **METHOD DETAILS**
  - Genotyping
  - Pathogen Load
  - Serum collection
  - IgG purification
  - IgG conjugation
  - Passive transfer of IgG
  - ELISA
  - Western blot for detection of αGal
  - Bacterial Strains and Culture Conditions
  - Flow cytometry of bacterial IgG binding and αGal expression
  - *In vivo* phagocytosis assay
  - Flow cytometry for bacterial staining
  - Flow cytometry for bacterial sorting
  - Extraction of bacterial DNA from feces
  - Extraction of sorted bacterial DNA
  - Metagenomics
  - Mathematical Modeling

- **QUANTIFICATION AND STATISTICAL ANALYSIS**
- **DATA AND SOFTWARE AVAILABILITY**

## SUPPLEMENTAL INFORMATION

Supplemental Information includes six figures and can be found with this article.

## CONTACT FOR REAGENTS AND RESOURCE SHARING

Further information and requests for reagents may be directed to, and will be fulfilled by, the lead contact, Miguel P. Soares.

## EXPERIMENTAL MODEL AND SUBJECT DETAILS

### Mice

Mice were used in accordance with protocols approved by the Ethics Committee of the Instituto Gulbenkian de Ciência (IGC) and Direção Geral de Alimentação e Veterinária (DGAV), following the Portuguese (Decreto-Lei no. 113/2013) and European (directive 2010/63/EU) legislation for animal housing, husbandry and welfare. C57BL/6J wild-type, *Ggta1*^*-/-*^ (Tearle et al., 1996), *J*_*h*_*t*^*-/-*^*Ggta1*^*-/-*^ (Gu et al., 1993), *Tcr*β^*-/-*^*Ggta1*^*-/-*^ (Yilmaz et al., 2014), *Aid*^*-/-*^*Ggta1*^*-/-*^ (Yilmaz et al., 2014) and *µs*^*-/-*^ *Ggta1*^*-/-*^ (Yilmaz et al., 2014) mice were used. *Iga*^*-/-*^*Ggta1*^*-/-*^ and *Rag2*^*-/-*^*Ggta1*^*-/-*^ mice were generated by crossing *Ggta1*^*-/-*^ (Tearle et al., 1996) with C57BL/6 *Iga*^*-/-*^ (Blutt et al., 2012) and *Rag2*^*-/-*^ (Shinkai et al., 1992) mice, respectively. Mice were bred and maintained under specific pathogen-free (SPF) conditions (12 h day/night, fed *ad libitum*), as described (Yilmaz et al., 2014).

Germ-free (GF) C57BL/6J *Ggta1*^+*/*+^ and *Ggta1*^*-/-*^ animals were bred and raised in the IGC gnotobiology facility in axenic isolators (La Calhene/ORM), as described (Yilmaz et al., 2014). Adult mice were transferred to sterile ISOcages (Tecniplast) and sterility of food, water, bedding, oral swab and feces were confirmed before each experiment by plating samples on Sabouraud Glucose Agar (BD #254039) for fungi, or Trypticase™Soy Agar II plates with 5% Sheep Blood (BD #254053) for bacteria, and incubated (37°C, 5 days) in air with 5% CO_2_ for aerobes and in air tight containers equipped with GasPak™ anaerobe container system (BD #11747194) for anaerobes. Anaerobic conditions were confirmed using BBL™ Dry Anaerobic Indicator Strips (Becton Dickinson #271051). Samples were added to Difco™ Nutrient Broth medium (#234000), incubated (37°C, 5 days) and plated (∼500 µL/plate) on Sabouraud Glucose Agar and Trypticase™Soy Agar II plates with 5% Sheep Blood and incubated (37°C, 5 days) under aerobic and anaerobic conditions. Plates and liquid medium were checked for absence of fungal and bacterial growth. Both male and female mice were used for all experiments. All animals were studied between 9-16 weeks of age unless otherwise indicated.

### Cecal Ligation and Puncture (CLP)

CLP was performed as described (Rittirsch et al., 2009). Procedures were performed routinely at the same time of the day, starting at 9 am. Briefly, mice were anaesthetized (intraperitoneal, *i*.*p*.) using ketamine (75 mg/kg) and xylazine (15 mg/kg) (∼140 µL/mouse, 1:1 vol/vol in sterile 0.9% saline). The lower left quadrant of the abdomen was disinfected with Betadine® solution. Under aseptic conditions, a 1- 2 cm lower left quadrant laparotomy was performed and the cecum with the adjoining intestine was exposed. 20-30% of the cecum below the ileo-cecal valve, was tightly ligated with a silk suture (3-0 Mersilk #W212) and perforated twice (23G needle). The cecum was then gently squeezed to extrude a small amount of cecal content from the perforation sites, returned to the peritoneal cavity and the abdomen was closed with silk sutures (Virgin Silk #C0761214). The skin was closed with Reflex 9 mm clips (Cellpoint Scientific #201-1000). Mice were resuscitated by injecting 800 µL (mice < 25 g) or 1 mL (mice > 25 g) of sterile 0.9% saline solution (subcutaneous, 25G needle). Mice were placed on a heating pad (30 min. - 2 h) until recovery from anesthesia and provided with free access to food and water by placing hydrogel or food pellets on the bottom of the cage. Mice were monitored every 12 h for survival for 14 days or euthanized at various time points for analysis of different parameters.

### Cecal Slurry Injection

Under aseptic conditions, 3-5 donor mice were sacrificed, and a 1-2 cm lower left quadrant laparotomy performed. The cecum was excised, contents extracted, pooled in pre-weighed sterile Eppendorf tubes and kept on ice. The cecal contents were weighed and homogenized in sterile PBS by vortexing (maximum speed, 1-5 min.). The resulting cecal slurry was filtered (BD Falcon™, 40 µm cell strainer, # 352340) and injected to recipient mice (*i*.*p*. 1-1.25 mg/g body weight, 25G needle). Mice were monitored every 12 h for survival for 14 days or euthanized at various time points for analysis of different parameters.

To analyze mouse survival when exposed to killed cecal content, cecal slurry from *Rag2*^*-/-*^*Ggta1*^*-/-*^ mice was prepared as above, pelleted by centrifugation (4,000 rpm, 4°C, 20 min.), supernatant discarded and material re-suspended in paraformaldehyde (PFA, 4% weight/vol in PBS). Fixation was left to proceed overnight, before centrifugation as above, and washing (2x, PBS). A lack of viable bacteria in the inoculum was confirmed by plating undiluted content on Trypticase™Soy Agar II plates with 5% Sheep Blood (BD #254053) and incubating at 37°C under anaerobic conditions for 3 days. Fixed cecal material was injected to mice (*i*.*p*. 1.25 mg/g body weight, 25G needle) and survival was monitored every 12 h for 14 days.

### Breeding experiments

Segregation of *Ggta1*^+*/*+^ and *Ggta1*^*-/-*^ genotypes carrying a similar microbiota derived from *Ggta1*^*-/-*^ mice was achieved, as described (Ubeda et al., 2012). Briefly, two or more breeding pairs were established, consisting of two *Ggta1*^*-/-*^ females and one *Ggta1*^+*/*+^ male per cage. The male was removed after one week and the females were placed in a clean cage until delivery. F_1_ *Ggta1*^+*/-*^ pups were weaned at 3-4 weeks of age and then co-housed until 8 weeks of age. F_1_ *Ggta1*^+*/-*^ breeding pairs were established randomly using one male and two females per cage. F_2_ pups were weaned at 3-4 weeks of age, genotyped and segregated according to their *Ggta1*^*-/-*^ *vs. Ggta1*^+*/*+^ genotype in separate cages until adulthood. Fecal pellets were collected (10-12 weeks of age) for microbiota analysis.

### Reproductive output

Breeding of *Ggta1*^+*/*+^ and *Ggta1*^*-/-*^ mice was performed under SPF conditions using trio breeding groups, composed of two females *per* male *per* cage. Breeding was established when mice reached 8-10 weeks of age. A total of n=432 of *Ggta1*^+*/*+^ and n=135 of *Ggta1*^*-/-*^ trio breeding groups were analyzed over a period of 5 years, spanning from 2012 to 2017. Breeding was monitored for: i) number of pups produced over the reproductive life-span of each breeding group, ii) number of pups per combined litter, whereby combined litter refers to the pool of offspring *per* breeding group, as accounted for at the time of weaning, iii) number of combined litters produced over the reproductive life-span of each breeding group and iv) reproductive senescence, as defined by the age at which females produced the last live combined litters. Breeding was followed until 2 months after the last viable litter. Pups were weaned at 3-4 weeks of age. Detection of dead progenitors and/or dead litters was a criterion for exclusion of the breeding group from the analysis.

## METHOD DETAILS

### Genotyping

Mice were genotyped from tail biopsies (0.5-1 cm) by PCR of genomic DNA using a standard protocol, *as per* manufacturer’s protocols (KAPA mouse genotyping kit #KK7352). Samples were lysed in KAPA Express Extract Enzyme (1 µL), KAPA Express Extract Buffer (5 µL) and water (44 µL), heated (75°C, 15 min., and 95°C, 5 min.), vortexed (3 sec.), centrifuged (16,000 g, 1 min., room temperature (RT)), diluted in water (1:4 vol/vol) and centrifuged (16,000 g, 1 min., RT). Extracted DNA (1 µL) was amplified in the PCR mix consisting of 2x KAPA2G Fast Genotyping Mix (5 µL), each primer (0.5 µL) and water (2-2.5 µL). Visualization of PCR products was done on a 1-2% agarose gel (80-100 V, 1-2 h).

### Pathogen Load

Mice were sacrificed 24 h after CLP or cecal slurry injection, placed in a sterile surgical field and sprayed thoroughly with ethanol. A 5×5 cm window was created on the abdomen by excising the skin. Ice-cold sterile PBS was injected (*i*.*p*. 5 mL, 25G needle). The mouse was shaken vigorously (5x horizontally and vertically) to homogenize the peritoneal fluid and the peritoneal lavage was collected (3-4 mL, 23G needle) and kept on ice. The abdominal and thoracic cavities were opened and blood was collected by intra-cardiac puncture through a 25G needle into a heparinized syringe. Mice were perfused (25 mL of ice-cold sterile PBS) through the left ventricle of the heart. The right atrium was cut after confirming blanching of the liver. Whole organs (*i*.*e*. lungs, liver, spleen and kidneys) were harvested, rinsed with sterile water and kept on ice in sterile Eppendorf tubes. Organs were homogenized under sterile conditions (2 mL dounce tissue grinder, Sigma #D8938-1SET) in 500 µL (lungs, kidneys and spleen) or 1 mL (liver) PBS. Serial dilutions were plated on Trypticase™Soy Agar II plates with 5% Sheep Blood (BD #254053) and incubated (37°C) in air with 5% CO_2_ for aerobes and in air tight containers equipped with GasPak™ anaerobe container system (Becton Dickinson #11747194) for anaerobes. Anaerobic conditions were confirmed using BBL™ Dry Anaerobic Indicator Strips (BD #271051). Colonies were counted after 24 h and quantified.

### Serum collection

Blood was collected from the submandibular vein of live mice (8-9 drops per mouse) or alternatively via intra-cardiac puncture of terminal mice. Coagulation was allowed to occur (1 h, RT), samples were centrifuged (2x, 2,000 g, 10 min., 4°C) and the supernatant was collected and stored at -20°C until use.

### IgG purification

IgG purification was performed using HiTrap™ Protein G HP (GE Healthcare #17-0404-01) according to manufacturer’s instructions, with modifications. Briefly, serum was pooled from 20-30 *Ggta1*^+*/*+^ or *Ggta1*^*-/-*^ mice, diluted in binding buffer (1:10 vol/vol, Tris 20 mM, pH 8.0, 150 mM NaCl) and filtered (PALL Lifesciences #4612, 0.2 µM). Samples were passed through the column (1 mL/min.) using a peristaltic pump (Gilson minipuls3) and the flow-through collected, followed by elution with elution buffer (100 mM glycine-HCl, pH 2.0, 1 mL/min.). The pH of the eluate (1 mL per Eppendorf tube) was neutralized (125 µL of Tris 1M, pH 9.0). The initial flow-through was passed through the column again and the process repeated whenever necessary. IgG concentration was initially measured by a spectrophotometer (NanoDrop™ 2000/2000C) and IgG fractions were pooled, dialyzed against PBS and concentrated (Amicon Ultra15 #UFC903024). Quality control was carried out by SDS-PAGE and the final IgG concentration was determined by ELISA, as described below.

### IgG conjugation

Purified mouse IgG was labeled with either PE/Cy5® (Abcam #ab102893) or AlexaFluor®594 (Molecular Probes #A10239), according to the manufacturer’s instructions. Concentration of the labeled IgG was determined by ELISA, as described below.

### Passive transfer of IgG

Purified mouse IgG (300 µg in 240 µL PBS) was injected (*i*.*p*.) to *Rag2*^*-/-*^*Ggta1*^*-/-*^ mice, 24 h before cecal slurry injection.

### ELISA

ELISA for IgG binding to cecal extracts was done, essentially as described (Zeng et al., 2016). Briefly, cecal content from 3-5 mice was collected, pooled and weighed in sterile pre-weighed Eppendorf tubes. The cecal content was then homogenized in sterile PBS by vortexing (maximum speed, 5 min., RT) and filtered (BD Falcon™, 40 µm cell strainer # 352340). Larger debris were removed by centrifugation (1000 rpm, 5 min., RT), the supernatant was collected and bacteria were pelleted by centrifugation (8,000 g, 5 min., RT). The pellets were washed in PBS (2x, 10,000 g, 1 min., RT) and re-suspended in PBS (500 µL). Bacteria were heat-killed (85°C, 1 h) and suspended in Carbonate-Bicarbonate buffer (0.5 M, pH 9.5, 50 µL/mg), producing a cecal lysate.

96-well ELISA plates (Nunc MaxiSorp #442404) were coated with the cecal lysate (100 µL/well, 4°C, overnight), washed (3x, PBS 0.05% Tween-20, Sigma-Aldrich #P7949-500ML), blocked (200 µL, PBS 1% BSA wt/vol, Calbiochem #12659-100GM, 3 h, RT) and washed (3x, PBS 0.05% Tween-20). Plates were incubated with serially diluted (50 µL) mouse sera (PBS 1% BSA, wt/vol), starting at 1:20 (vol/vol) for detection of IgG1, IgG2c and IgG3 and at 1:200 (vol/vol) for detection of IgG2b (2 h, RT) and washed (5x, PBS/0.05% Tween-20). Immunoglobulins were detected using horseradish peroxidase (HRP)-conjugated goat anti-mouse IgG1 (Southern Biotech #1070-05), IgG2b (Southern Biotech #1090-05), IgG2c (Southern Biotech #1079-05) or IgG3 (Southern Biotech #1100-05) in PBS/1%BSA/0.01% Tween-20 (100 µL, 1:4,000 vol/vol, 1 h, RT) and plates were washed (5x, PBS/0.05% Tween-20).

For quantification of total serum IgG, 96-well ELISA plates (Nunc MaxiSorp #442404) were coated with goat anti-mouse IgG (Southern Biotech #1030-01, 2 µg/mL in Carbonate-Bicarbonate buffer, 100 µL/well, overnight, 4°C), washed (3x, PBS 0.05% Tween-20, Sigma-Aldrich #P7949-500ML), blocked (200 µL, PBS 1% BSA wt/vol, 2 h, RT) and washed (3x, PBS 0.05% Tween-20). Plates were incubated with serially diluted mouse sera (50 µL, PBS 1% BSA, wt/vol, 2 h, RT) and standard mouse IgG (Southern Biotech #0107-10, prepared in duplicates) and washed (5x, PBS/0.05% Tween-20). IgG was detected using HRP-conjugated goat anti-mouse IgG (Southern Biotech #1030-05) in PBS/1%BSA/0.01% Tween-20 (100 µL, 1:4,000 vol/vol, 1 h, RT) and plates were washed (5x, PBS/0.05% Tween-20).

For quantification of anti-αGal IgG, 96-well ELISA plates (Nunc MaxiSorp #442404) were coated with either αGal-BSA (Dextra) or goat anti-mouse Ig(H+L) (Southern Biotech #1010-01) (5 µg/mL in Carbonate-Bicarbonate buffer, 50 µL/well, overnight, 4°C). Wells were washed (3x, PBS 0.05% Tween-20, Sigma-Aldrich #P7949-500ML), blocked (200 µL, PBS 1% BSA wt/vol, 2 h, RT) and washed (3x, PBS 0.05% Tween-20). Plates were incubated with serially diluted IgG purified from *Ggta1*^+*/*+^ or *Ggta1*^*-/-*^ mice (50 µL, PBS 1% BSA, wt/vol, 2 h, RT) and standard mouse anti-αGal IgG2b (derived from GT6-27 (Ding et al., 2008) and washed (5x, PBS/0.05% Tween-20). IgG was detected using HRP-conjugated goat anti-mouse IgG2b (Southern Biotech #1090-05) in PBS/1%BSA (100 µL, 1:4000 vol/vol, 1.5 h, RT) and plates were washed (5x, PBS/0.05% Tween-20).

For IgG binding to FcγRs, 96-well ELISA plates (Nunc MaxiSorp #442404) were coated with purified IgG (5 µg/mL in Carbonate-Bicarbonate buffer, 50 µL/well, overnight, 4°C), washed (3x, PBS 0.05% Tween-20, Sigma-Aldrich #P7949-500ML), blocked (200 µL, PBS 1% BSA wt/vol, 2 h, RT) and washed (3x, PBS 0.05% Tween-20). Plates were incubated with serially diluted biotinylated mouse FcγRI (Acrobiosystems #CD4-M5227), FcγRIIb (Acrobiosystems #CDB-M82E8), FcγRIII (Acrobiosystems #CDA-M52H8), FcγRIV (Acrobiosystems #FC4-M82E8) or FcRn (Acrobiosystems #FCM-M82W5) (50 µL, PBS 1% BSA, wt/vol, 2 h, RT) and washed (5x, PBS/0.05% Tween-20). Signal was detected using HRP-conjugated Streptavidin in PBS/1%BSA/0.01% Tween-20 (100 µL, 1:2,000 vol/vol, 1 h, RT) and plates were washed (5x, PBS/0.05% Tween-20).

For IgG binding to C1q, 96-well ELISA plates (Nunc MaxiSorp #442404) were coated with purified IgG (5 µg/mL in Carbonate-Bicarbonate buffer, 50 µL/well, overnight, 4°C), washed (3x, PBS 0.05% Tween-20, Sigma-Aldrich #P7949-500ML), blocked (200 µL, PBS 1% BSA wt/vol, 2 h, RT) and washed (3x, PBS 0.05% Tween-20). Plates were incubated with serially diluted mouse C1q (CompTech #M099), washed (5x, PBS/0.05% Tween-20), incubated with biotinylated anti-mouse C1q (Cedarlane #CL7501B, 1:50,000 vol/vol, PBS/1%BSA/0.01% Tween-20, 1 h, RT) and washed (5x, PBS/0.05% Tween-20). Signal was detected using HRP-conjugated Streptavidin in PBS/1%BSA/0.01% Tween-20 (100 µL, 1:2,000 vol/vol, 1 h, RT) and plates were washed (5x, PBS/0.05% Tween-20).

To control for IgG binding, 96-well ELISA plates (Nunc MaxiSorp #442404) were coated with purified IgG (5 µg/mL in Carbonate-Bicarbonate buffer, 50 µL/well, overnight, 4°C), washed (3x, PBS 0.05% Tween-20, Sigma-Aldrich #P7949-500ML), blocked (200 µL, PBS 1% BSA wt/vol, 2 h, RT) and washed (3x, PBS 0.05% Tween-20). Plates were incubated with serially diluted HRP-conjugated goat anti-mouse IgG (Southern Biotech #1030-05) in PBS/1%BSA/0.01% Tween-20 (100 µL, 1 h, RT) and washed (5x, PBS/0.05% Tween-20).

HRP activity was detected with 3,3′,5,5′-Tetramethylbenzidine (TMB) Substrate Reagent (BD Biosciences #555214, 50 µL, 20-25 min., dark, RT) and the reaction was stopped using sulfuric acid (2N, 50 µL). Optical densities (OD) were measured using a MultiSkan Go spectrophotometer (ThermoFisher) at *λ*=450 nm and normalized by subtracting background OD values (*λ*= 600 nm). Concentrations of IgG2b in purified serum IgG samples were calculated from the absorbance obtained with reference to the standard curve determined for total and αGal-specific IgG2b, respectively.

### Western blot for detection of αGal

Purified IgG, BSA conjugated to αGal (Dextra # NGP1334) and unconjugated BSA (New England Biolabs #174B9000S) were denatured in Laemmli buffer (1% β-mercaptoethanol, 2% SDS, 70°C, 10 min.) and separated by SDS-PAGE (12% acrylamide/bisacrylamide gel, 29:1; 100V; 2 h). Proteins were transferred onto a PVDF membrane (50 min, 12 V), blocked (5% BSA in 20 mM Tris/HCl pH 7.5, 150 mM NaCl and 0.1% Tween-20 or TBST buffer, 2 h) and incubated (4°C, overnight) with biotinylated BSI-B4 lectin from *Bandeiraea* (*Griffonia) simplicifolia* (Sigma-Aldrich, #L3759-1MG, 1 mg/mL, 5% BSA in TBST buffer, 5 mL) or with Anti-αGal M86 mAb (1:1000, 5% BSA in TBST buffer, 5 mL). Membranes were washed with TBST (1x, 5 min., RT) and incubated with Streptavidin-HRP (10 mL, 1:5,000, 1 h, RT) for detection of BSI-B4, or with goat anti-mouse IgM-HRP (Southern Biotech #1021-05, 10 mL, 1:5,000, 1 h, RT) for detection of M86 mAb. Membranes were washed with TBST (3x, 20 min., RT) and developed (SuperSignal® West FEMTO Max. Sensitivity Substrate #11859290). As a loading control, SDS-PAGE gel was stained with InstantBlue™ Safe Coomassie Stain (Sigma # ISB1L-1L).

### Bacterial Strains and Culture Conditions

*E. coli* O86:B7 (ATCC12701) were streaked from -80°C glycerol stocks onto Luria-Bertani (LB) 1.5% agar plates, incubated at 37°C overnight. For liquid culture, a single colony was inoculated into 5-10 mL LB liquid medium and incubated (12-16 h, 37°C) with aeration (180-200 rpm). For analysis of αGal expression, *E. coli* O86:B7 was grown in NB medium (BD Difco). Optical Density at 600 nm (OD_600_) of the overnight culture was measured by spectrophotometry (Bio-Rad SmartSpec™3000). To harvest bacteria during exponential growth phase, sub-culturing was done in NB medium with a starting OD_600_ of 0.05 by incubation (3 h, 37°C) with aeration (180-200 rpm). OD_600_ of the bacterial sub-culture was measured (OD=2 corresponding to approximately 10^9^ CFU/mL). The culture was incubated to stop growth (5 min., 4°C) and bacteria were pelleted by centrifugation (4,000 rpm, 20 min., 4°C). The pellet was suspended in sterile PBS (5 mL). OD_600_ was measured in PBS (1:10 vol/vol) in triplicate. A volume of bacterial culture corresponding to approximately 10^6^-10^7^ cells per condition was harvested, fixed with 4% PFA/1xPBS and stained for flow cytometry as described.

*E. coli* M6L4, *E. coli* M5S5 and *Klebsiella pneumoniae* were cultured as described above. Cells from -80°C frozen stocks were streaked onto LB agar plates, incubated overnight at 37°C under aerobic conditions, and cells from the resulting colonies used to inoculate 5-10 mL LB medium. Bacterial cultures were incubated at 37°C, with aeration (180-200 rpm), for 12-16 h. *Staphylococcus sciuri, Clostridium bifermentans, Enterococcus faecalis* and *E. gallinarum* were grown as above using solid or liquid brain heart infusion (BHI) media supplemented with 100 mM vitamin K1 and 1.9 μM hematin; and cultured anaerobically at 37°C for 2 days. *E. faecalis* and *E. gallinarum* were also grown under aerobic conditions on BHI solid or liquid media overnight for flow cytometry analysis. *Parabacteroides goldsteinii* was cultured anaerobically on solid or liquid BHI medium supplemented with 1.2 µM histidine, 1.9 µM hematin, 1 µg/mL menadione, and 500 µg/mL cysteine at 37°C for 2 days.

### Flow cytometry of bacterial IgG binding and αGal expression

Overnight cultures of bacteria were prepared as described above. Samples of 5-20 µL of each bacterial culture depending on OD_600_ measurements, and corresponding to approximately 10^6^-10^7^ cells, were fixed in PFA (4% wt/vol in PBS) and washed with filter-sterilized PBS. Bacteria were incubated in either Fluorescein Isothiocyanate (FITC)-conjugated BSI-B4 lectin from *Bandeiraea* (*Griffonia) simplicifolia* (Sigma-Aldrich, #L2895-1MG, 50 µL, 40 µg/mL in PBS, 2 h) for detection of αGal, or IgG purified from *Ggta1*^+*/*+^ or *Ggta1*^*-/-*^ mouse serum (60 µg/mL, PBS 2% BSA, wt/vol, 30 min.) followed by anti-mouse IgG-PE (Southern Biotech #1030-09, 1:100 in PBS 2% BSA wt/vol, 30 min.). Cells were washed and analyzed by flow cytometry using an LSR Fortessa SORP (BD Biosciences), as described below. Data from at least 10,000 single cell events were measured and analyzed using FlowJo software (v.10).

### *In vivo* phagocytosis assay

*In vivo* phagocytosis assays were performed upon injection (i.p.) of FITC-labeled *E. faecalis* (10^8^ CFU, 4% PFA, in 100 μL PBS) to *J*_*h*_*t*^*-/-*^*Ggta1*^*-/-*^ mice. Bacteria were either unopsonized or opsonized with IgG (300 μg/mL, 30 min., 37°C) purified from *Ggta1*^+*/*+^ or *Ggta1*^*-/-*^ mice. Peritoneal lavage was done 3 h after injection (5 mL, ice-cold PBS) using naïve mice as controls. Samples were prepared as follows: 500 μL aliquots of the lavage were centrifuged (500 g, 2 min., 4°C) and viable cells were stained using Live/dead fixable yellow stain (1:1000 in 1% FBS/PBS, Molecular Probes, 15 min., on ice). Cells were washed and incubated (15-20 min., on ice) with Fc Block (1:100 in 1% FBS/PBS, Clone 2.4G2, BD Biosciences), followed by anti-CD45-PE-Cy5.5 (1:100, clone 30-F11, Life Technologies), anti-Ly6G-PE-Cy7 (1:100, clone 1A8, Biolegend) and anti-CD11b-BV421 (1:100, clone M1/70, Biolegend). Cells were analyzed using LSR Fortessa X20 (BD Biosciences) and FACSDiva software (BDv.6.2). Cell numbers were calculated using PerfectCount Microspheres (Cytognos). Data from at least 10,000 single viable CD45^+^ cells were acquired and analyzed by FlowJo software (v10.0.7).

### Flow cytometry for bacterial staining

Bacterial staining of cecal content was performed essentially as described (Bunker et al., 2017; Koch et al., 2016; Zeng et al., 2016). Briefly, cecal slurry was prepared as described above, homogenized, filtered through a 40 μM cell strainer and diluted to a concentration of 5 mg/mL in sterile PBS. Large debris were pelleted by centrifugation (600 g, 4°C, 5 min.). 50 μL supernatant (containing bacteria) per condition was added to a 96-well v-bottom plate (Corning Costar #3894) for staining. Bacteria were pelleted by centrifugation (3,700 g, 10 min., 4°C) and suspended in flow cytometry buffer (filter-sterilized 1xPBS, 2% BSA, wt/vol). Bacterial DNA was stained using SYTO®41 Blue Fluorescent Nucleic Acid Stain (Molecular Probes #S11352, 1:200 vol/vol, wt/vol) or with SYBR® Green I Nucleic Acid Gel Stain (Invitrogen #S-7563, 1:1000 vol/vol) in flow cytometry buffer (100 µL, 30 min., RT). Cecal content from germ free (GF) mice was used as control. Bacteria were washed in flow cytometry buffer (200 µL), centrifuged (4000 g, 10 min., 4°C) and supernatant was removed by flicking the plate. Bacteria were incubated with mouse serum (1:20, vol/vol), or purified IgG from *Ggta1*^+*/*+^ or *Ggta1*^*-/-*^ mice (60 µg/mL) in flow cytometry buffer (100 µL, 30 min., RT) and washed, as above. IgG was detected using Phycoerythrin (PE)-labeled goat anti-mouse IgG (Southern Biotech #1030-09, 100 µL, 1:100 vol/vol, 30 min., RT or 4°C, 1 h) and washed as above. Samples were re-suspended in flow cytometry buffer (300 µL), transferred to round-bottom tubes (BD Falcon™ #352235) and centrifuged (300 g, 1 min., RT).

For co-staining of cecal content with purified mouse IgG from both genotypes, the bacteria were collected, stained for DNA and washed, as described above. Bacteria were incubated with Alexa-Fluor 594 (AF594)-conjugated mouse IgG and PeCy5-conjugated mouse IgG (100 µL, 60 µg/mL, 30 min., RT). Samples were washed and collected before analysis, as described above.

For co-staining of cecal content with purified mouse IgG and αGal, bacteria from the cecal content were collected, stained for DNA and washed, as described above. Bacteria were incubated in Fluorescein Isothiocyanate (FITC)-conjugated BSI-B4 lectin from *Bandeiraea* (*Griffonia) simplicifolia* (Sigma-Aldrich, #L2895-1MG, 50 µL, 40 µg/mL in PBS, 2 h) for detection of αGal and IgG purified from *Ggta1*^+*/*+^ or *Ggta1*^*-/-*^ mouse serum (60 µg/mL, PBS 2% BSA, wt/vol, 30 min.) followed by anti-mouse IgG-PE (Southern Biotech #1030-09, 1:100 in PBS 2% BSA wt/vol, 30 min.). Supernatant was removed by aspiration and the stained bacteria were suspended in sterile filtered PBS (500 µL) and collected before analysis as described above. *E coli* O86:B7 (about 10^7^ per tube) were also stained with FITC-conjugated BSI-B4 lectin as described above, as a positive control for bacterial αGal expression.

For bacterial staining in the peritoneal cavity, the bacteria were harvested by peritoneal lavage as described above, and centrifuged (900 *g*, 5 min., 4°C). The supernatant containing the bacteria was collected (500 µL), centrifuged (10,000 g, 1 min., 4°C) and suspended in flow cytometry buffer. The remainder of the procedure was similar to that detailed above. An additional step to exclude host leukocytes was included in which the suspension was co-stained with PECy5.5-labeled rat anti-mouse CD45 (eBioscience, Clone 30-F11, 1:100 vol/vol, 15 min., RT or 4°C, 1 h) and washed as above. For staining αGal, the pellet was suspended in freshly prepared BSI-B4-FITC conjugate (50 µL, 40 µg/mL, 2 h, RT). Samples were washed and collected before analysis as described above.

Samples were analyzed in LSR Fortessa SORP (BD Biosciences) equipped with a high-throughput sampler (HTS) using the FACSDiva Software (BD v.6.2). SYBR® Green and BSI-B4-FITC were excited with blue laser (488 nm, filters: 530/30, 502LP), IgG-PE with Yellow-green laser (561 nm, detection filter: 590/20), CD45-PECy5.5 with Yellow-green laser (561 nm, detection filters: 705/70, 685LP), Syto41 with Blue-violet laser (442 nm, detection filters: 480/40, 455LP), PeCy5 with Blue laser (488 nm, detection filters: 695/40, 660LP) and AF594 with Yellow-green laser (561 nm, detection filters: 630/30, 610LP). Compensations were done using anti-rat/hamster Igκ compensation beads (BD™ CompBead #552845). Before acquisition of samples, laser voltages were standardized using SPHERO™ Ultra Rainbow Fluorescent Particles (Spherotech #URFP01-30-2K). Data from at least 10,000 single bacterial events were measured and analyzed by FlowJo software (v10.0.7).

### Flow cytometry for bacterial sorting

Bacterial staining for IgG in the peritoneal cavity was done as described above. Sorting was performed in FACSAria IIu (BD Biosciences, 70 µm nozzle). SYBR® Green, IgG-PE and CD45-PE-Cy5.5 were excited with blue laser (488 nm) and fluorescence detected using the following filters, respectively: 530/30, 502LP, 585/42, 550LP, 695/40, 655LP. The gating strategy was set using Fluorescence Minus One (FMO) controls for all fluorochromes, as well as biological controls that specifically lack target populations. 5,000-10,000 events of IgG-positive and IgG-negative bacterial populations were sorted into tubes containing filter-sterilized PBS (50 µL) and stored at -80°C until further analysis.

### Extraction of bacterial DNA from feces

Fecal pellets (4-5/mouse) were collected in sterile Eppendorfs and stored at -80°C. DNA extraction was done according to manufacturer’s instructions (QIAamp Fast DNA Stool Mini Kit #50951604). Briefly, individual samples were thawed and mechanically disrupted in InhibitEX Buffer (1 mL) with a motorized pestle using sterile glass beads (Disrupter Genie # 9730100, about 0.4 g/sample). Disruption of microbial cells was enhanced using TissueLyser II (QIAGEN, 30 shakes/sec., 1 min., 2x). To lyse Gram-negative bacteria, the suspensions were heated (12 min., 95°C), vortexed (15 s) and larger particles were pelleted by centrifugation (20,000 g, 1 min., RT). The supernatant (200 µL) was lysed by incubation (12 min., 70°C) in a mixture of Proteinase K (15 µL) and AL buffer (200 µL) and vortexed (15 s). DNA in the lysate was precipitated with ethanol (96–100%, 200 µL) and washed in AW buffer (20,000 g, 1min., RT, 2x). DNA was incubated in ATE buffer (100 µL, 1 min., RT), eluted by centrifugation (20,000 g, 1 min., RT) and frozen at -80°C until further use.

### Extraction of sorted bacterial DNA

DNA was extracted from flow cytometry sorted bacterial samples using manufacturer’s protocols (DNeasy PowerSoil Kit #12888-50). Briefly, samples (5×10^3^-1×10^4^ bacteria) were lysed in Solution C1 (60 µL) in PowerBead tubes, vortexed briefly and heated (10 min., 65°C). Mechanical disruption of microbial cells was done using TissueLyser II (QIAGEN, 30 shakes/sec., 10-20 min.) and the samples were centrifuged (10,000 *g*, 30 sec., RT). The supernatant was cleaned by incubation (5 min., 2–8°C) in Solution C2 (250 µL) followed by Solution C3 (200 µL), vortexed (5 sec.), and centrifuged (10,000 *g*, 1 min., RT). To bind DNA to the MB Spin Column, the supernatant was mixed with Solution C4 (1,000 µL), vortexed (5 sec.), loaded onto the column and centrifuged (10,000 *g*, 1 min., RT). DNA was washed in Solution C5 (500 µL) by centrifugation (10,000 *g*, 30 sec., RT, 2x). DNA was eluted in Solution C6 (50 µL) by centrifugation (10,000 *g*, 30 sec., RT) and stored at -80°C until further use.

### Metagenomics: 16S amplicons production and sequencing

The 16S rRNA V4 region was amplified in triplicate by PCR following the Earth Microbiome Project (http://www.earthmicrobiome.org/emp-standard-protocols/). Briefly, the mix for each sample consisted of DNA (1 µL), water (9 µL), PCR Mastermix (5PRIME HotMasterMix-1000R #733-2474, 2x, 10 µL,), Primer_barcode (2 µM, 2.5 µL) and Primer_universal (2 µM, 2.5 µL). For samples extracted from bacterial sorting (low biomass), 10 µL of DNA and no water were used. The PCR conditions were: 94°C (3 min.); 35 cycles of 94°C (45 sec.), 50°C (60 sec.), 72°C (90 sec.); 72°C (10 min.), 4°C (1 h) for 96 wells and 94°C (3 min.); 35 cycles of 94°C (60 sec.), 50°C (60 sec.), 72°C (105 sec.); 72°C (10 min.), 4°C (1 h). After amplification, the triplicates of each sample were pooled (75 µL) and quantified with Quant-iT™ PicoGreen® dsDNA Assay Kit (Invitrogen™#P7589). Equal amounts of amplicons from each sample containing individual barcodes were pooled (240-300 ng for low biomass) and cleaned (MoBio UltraClean PCR Clean-Up Kit #12500). DNA purity and concentration were assessed using a spectrophotometer (NanoDrop™) and Qubit® dsDNA HS Assay Kit (Invitrogen™) #Q32854). DNA library was prepared by denaturing with NaOH (0.2 M, 5 min., RT) and mixed with Illumina Phix (10-15%) to balance the nucleotide representation.

Sequencing of the 16S rRNA region was done using custom primers from the Earth Microbiome Project that was adapted for the Illumina MiSeq (MiSeq Reagent Kit v2, 500 cycle #MS-102-2003, Illumina) (Caporaso et al., 2010b; Zhang et al., 2014). The custom primers were: Read 1 primer (TATGGTAATTGTGTGYCAGCMGCCGCGGTAA), Read 2 primer (AGTCAGCCAGCCGGACTACNVGGGTWTCTAAT) and Index primer (AATGATACGGCGACCACCGAGATCTACACGCT). The denatured DNA library and the custom primers were loaded on specific reservoirs on the MiSeq cartridge and sequenced on a 2×250 cycles run.

### Metagenomics: Sequences and Operational Taxonomic Unit (OTU) Quality Control

The raw sequencing reads obtained from Illumina MiSeq were demultiplexed and quality filtered with SeqTK (v.1.2-r94) using q=0.01. Filtered paired reads were merged with PEAR (v.0.9.6) (Zhang et al., 2014) using default parameters. Merged reads were then processed using the QIIME package (v.1.9.1) (Caporaso et al., 2010b). Reads included had a quality score above 19, a median length of 250 bp, maximum 1 mismatch on the primer and default values for other quality parameters. Quality filtered reads were clustered into OTUs at 97% similarity using the default UCLUST (Edgar, 2010) method with an open reference approach. Subsequent taxonomic assignment was performed using the UCLUST classifier against the Greengenes database (v.13.8) (DeSantis et al., 2006). Sequence alignment and open-reference OTU picking (Rideout et al., 2014) were performed using the default Pynast (Caporaso et al., 2010a). Tree building was done with (Price et al., 2010) and taxonomic assignment with the RDP classifier (Wang et al., 2007). Two extra filtering steps were applied to taxonomy-assigned OTUs to remove outliers, eliminating sequences with less than a total of 10 counts across all samples and sequences with more than 50 counts for 3 samples or less.

### Metagenomics: Downstream Bioinformatics Analyses

To calculate alpha-diversity measures, samples were rarefied to match the least abundant sample. Using QIIME 1.9.1 (Caporaso et al., 2010b), the Chao1 and Shannon diversity measures were obtained for each sample and the mean of 10 independent rarefactions was taken. To estimate the significance of differences of alpha-diversity, the Mann-Whitney test was used to compare two groups and Kruskall-Wallis to compare multiple groups followed by Dunn post hoc test (Dunn, 1964) and Bonferroni correction was done to estimate significance of pairwise differences. The Unweighted and Weighted UniFrac distances (Lozupone et al., 2011) between samples were calculated, Principal Coordinates Analysis (PCoA) was performed and group significance estimated by using PERMANOVA test to obtain a pseudo-F statistic and a p-value for the statistic. In the case of multivalued factors, PERMANOVA was executed on all pairwise comparisons, followed by Bonferroni correction. Alternatively, the Mann-Whitney or Kruskall-Wallis test were used to compare mean Unweighted and Weighted UniFrac distances between elements of different groups. Ellipses are centered on the categorical averages of the metric distances with a 95% confidence interval for the first two coordinates of each group, drawn on the associated PCoA.

Linear discriminant analysis effect size (LEfSe) analysis (Segata et al., 2011) was performed to represent taxa distinguishing different groups. Cladograms, generated from LEfSe analysis, represent taxa enriched in each genotype. The central point represents the root of the tree (Bacteria), and each ring represents the next lower taxonomic level (phylum through genus). The diameter of each circle represents the relative abundance of the taxon.

### Mathematical Modeling: Model structure and assumptions

The model computes the lifetime reproductive success of a typical host from a given genotype. To obtain the reproductive output over lifetime, the birth rate at each age, *B(a)*, was weighted by the survival probability, *S(a)*, at that age, and integrated over the entire lifespan. Denoting this lifetime reproductive success as fitness *W*, this quantity was formalized as follows:

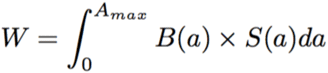

where *A*_*max*_ is the maximum lifespan of a typical host. In the absence of infection, the survival probability over ages follows a simple exponential with rate equal to the natural host mortality rate *(m)*:

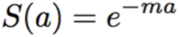

However, in the presence of exposure to infection, the survival probability becomes a weighted average between probability of getting infected/exposed *(E)* and subsequently dying from infection, and the probability of escaping infection *(1*−*E)* and dying from natural mortality. The parameter *E* denoting the lifetime infection risk of 0≤E≤1, constant over age, is assumed equal for both genotypes and driven by the environment. Its influence on the survival function of a typical host from the first genotype was formalized as follows:

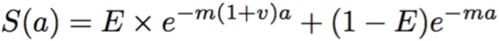

where *v* is the fold-increase in mortality due to infection, an indicator of pathogen virulence. For example if *v = 5*, a typical infected host will experience a 5-fold additional increase in mortality relative to an uninfected host per unit of time.

Protection of a second host genotype from infection was modeled through a factor *q (0* ≤ *q < 1)*, relative to the reference genotype, reducing the fraction of hosts that experience infection-induced mortality. Motivated by the data, it was assumed that individuals dying of infection experience the same mortality rate. Thus, the age-specific survival function of a typical host from the second genotype is given by:

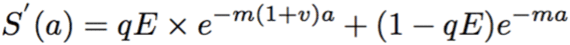

The protective effect is defined as *p = 1* − *q*, namely the relative gain in proportion survival of infected hosts in the second genotype relative to the first. When comparing the two host genotypes (0/1) in terms of their lifetime reproductive success, their difference in birth rate given by *B*_*0*_*(a)* for host genotype 0 and *B*_*1*_*(a)* for host genotype 1, was combined with their difference in survival given by *S*_*0*_*(a)* and *S*_*1*_*(a)*, accounting for infection. This led to the numerical comparison of *W*_*0*_ and *W*_*1*_:

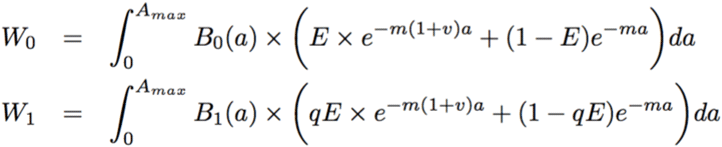

With this model for lifetime reproductive output, the fitness ratio was given by:

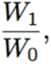

which if above 1, indicates a relative advantage of host genotype 0, and if below 1, indicates an advantage of host genotype 1. The selective advantage of 1 to 0 was then defined as:

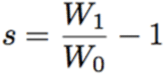

If s is positive (s > 0), the higher it will be, the more host genotype 1 are favored by natural selection relative to host genotype 0. Conversely, if s is negative (s < 0), then host genotype 1 suffer overall a fitness disadvantage relative to host genotype 0. Variation in model parameters common to or different between the two host genotypes in corresponding survival/birth rate functions, leads to explicit mathematical variation in *s*, and provides insights on biological conditions favoring selection.

### Mathematical Modeling: Model application to the dataset

We numerically parameterized the model based on our study with laboratory mice, taking *Ggta1*^+*/*+^ as reference host genotype 0 and *Ggta1*^*-/-*^ as host genotype 1. Natural lifespan was assumed to be 128 weeks (Kunstyr and Leuenberger, 1975), leading to a natural mortality rate *m=0*.*0078* per week. The survival data from the CLP experiments with the two genotypes (*Fig. 1A*) were used for the estimation of *v* (the fold-increase in mortality rate due to infection) and *p* (relative reduction in fraction of *Ggta1*^*-/-*^ hosts that die of infection). Since all mortality occurred within 2 weeks, it was assumed that this corresponds to all mortality due to infection in each group, (1-0.676) for *Ggta1*^*-/-*^, and (1-0.089) for *Ggta1*^+*/*+^, leading to a relative protective effect of 64% (*p=0*.*64*). The infection-induced mortality via an additional factor *v* relative to natural mortality, was calculated as:

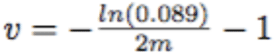

which in our particular case resulted in *v=154*, indicating substantial virulence. Our data support similar death kinetics in both genotypes, motivating the same *v* parameter in their survival functions. While capturing the difference between genotypes with a single parameter *p* is appropriate and sufficient in our context, in other systems, *v* could also vary. Exposure to infection *E* was assumed to be constant over age in this model. However, an age-dependent exposure *E(a)* could also be used, informed by empirical evidence or theoretical assumptions. Increasing exposure risk differentially in younger ages should amplify the selection potential for protection against infection. In contrast, increasing exposure risk in the post-reproductive period should reduce the selection potential for protection, given that host fitness would be more strongly affected by the reproductive fitness cost in that case. In the current formalism, epidemiological and co-evolutionary loops between host and pathogens were not modeled. The source of infection was assumed to be environmental and not explicitly driven by transmission between hosts. Similarly, details of pathogen-immunity dynamics within the host were not included. More complex modeling frameworks capturing such nested and bidirectional population feedbacks (Gilchrist and Sasaki, 2002), were considered to be beyond the scope of the current work.

### Mathematical Modeling: quantification and statistical analyzes

All statistical tests were performed using GraphPad Prism Software (v.6.0). To assess differences in binding of purified IgG to each of the Fcγ receptors and C1q, the sigmoidal curves of the form:

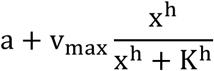

where x is the concentration, to each of the OD curves, were fitted using least-squares regression. One initial fit to the aggregate data was used to provide initial estimates to the fitting algorithm. Estimates for K (concentration at which OD is half-maximum) were compared between *Ggta1*^+*/*+^ and *Ggta1*^*-/-*^ IgG preparations using Mann-Whitney tests for differences in the median. Curve fitting and statistical tests were performed using scipy 1.2.1.

## SUPPLEMENTAL FIGURE TITLES AND LEGENDS

**Figure S1. *Ggta1* and *Rag2* deletion are associated with changes in microbiota composition. Related to Figure 1.**

**A-B**) Microbiota **A**) richness (Chao index) and **B**) diversity (Shannon index) in the same samples as in *Fig. 1C-E*. **C-D**) Microbiota **C**) richness (Chao index) and **D**) diversity (Shannon index) in the same samples as in *Fig. 1G-H*. **E**) Breeding strategy for the generation of *Rag2*^*-/-*^*Ggta1*^*-/-*^ mice from *Ggta1*^*-/-*^ females. F_0_ *Rag2*^*-/-*^*Ggta1*^+*/*+^ males crossed with *Rag2*^+*/*+^*Ggta1*^*-/-*^ females to generate F_1_ *Rag2*^+*/-*^*Ggta1*^+*/-*^ mice, which were bred to generate F_2_ *Rag2*^*-/-*^*Ggta1*^*-/-*^ *vs. Rag2*^+*/*+^*Ggta1*^*-/-*^ littermates. F_2_ *Rag2*^*-/-*^*Ggta1*^*-/-*^ mice were bred subsequently for 7 generations. **F-I**) Microbiota PCoA of **F**) Unweighted UniFrac, **G**) Weighted UniFrac distance, **H**) richness (Chao index) and **I**) diversity (Shannon index) from 16S rRNA amplicons in fecal samples from *Rag2*^+*/*+^*Ggta1*^*-/-*^ (n=15) and *Rag2*^*-/-*^*Ggta1*^*-/-*^ (n=14) mice. **J**) LDA scores and **K**) Cladogram, generated from LEfSe analysis, showing taxa enriched in *Rag2*^+*/*+^*Ggta1*^*-/-*^ (red) *vs. Rag2*^*-/-*^*Ggta1*^*-/-*^ (green) fecal microbiota in the same samples as (F-I); a: family_Prevotellaceae, b: family_Rikenellaceae, c: family_24-7, d: class_Alphaproteobacteria, e: class_Betaproteobacteria, f: order_RF32, g: order_Burkholderiales, h: order_RF39, i: order_Anaeroplasmatales. Data from one experiment. **L**) Survival of *Ggta1*^+*/*+^ (n=5) and *Ggta1*^*-/-*^ (n=4) mice infected (*i*.*p*.) with paraformaldehyde-treated cecal content from *Rag2*^*-/-*^*Ggta1*^*-/-*^ mice. Data from one experiment. Symbols in (A, B, C, D, F, G, H, I) are individual mice. Red bars (A, B, C, D, H, I) correspond to mean values. Error bars (A, B, C, D, H, I) correspond to SD. P values in (A, B, C, D, H, I) are calculated using Mann-Whitney test, P values and Fs in (F, G) using PERMANOVA test and in (L) using log-rank test. *P < 0.05, **P < 0.01, ***P < 0.005; ns: not significant

**Figure S2. Loss of *Ggta1* enhances resistance to systemic polymicrobial infection via a mechanism independent of IgM, IgA, or T cells. Related to Figure 2.**

**A**) Survival of *µS*^+*/*+^*Ggta1*^*-/-*^ (n=6) and *µS*^*-/-*^*Ggta1*^*-/-*^ (n=10) mice infected (*i*.*p*.) with a cecal inoculum from *Rag2*^*-/-*^*Ggta1*^*-/-*^ mice; 2 experiments. **B**) Colony forming units (CFU) of aerobic (Ae) and anaerobic (An) bacteria of *µS*^+*/*+^*Ggta1*^*-/-*^ (n=5) and *µS*^*-/-*^ *Ggta1*^*-/-*^ (n=4-5) mice, 24 hours after infection; 2 experiments. **C**) Survival of *Iga*^+*/*+^*Ggta1*^*-/-*^ (n=9) and *Iga*^*-/-*^*Ggta1*^*-/-*^ (n=11) mice infected as in (A); 2 experiments. **D**) CFU of Ae and An bacteria of *Iga*^+*/*+^*Ggta1*^*-/-*^ (n=5) and *Iga*^*-/-*^*Ggta1*^*-/-*^ (n=6) mice, 24 hours after infection; 2 experiments. **E**) Survival of *Tcrβ*^+*/*+^*Ggta1*^*-/-*^ (n=7) and *Tcrβ*^*-/-*^ *Ggta1*^*-/-*^ (n=8) mice infected as in (A); 2 experiments. **F**) CFU of Ae and An bacteria of *Tcrβ*^+*/*+^*Ggta1*^*-/-*^ (n=3) and *Tcrβ*^*-/-*^*Ggta1*^*-/-*^ (n=6) mice, 24 hours after infection; 2 experiments. **G**) Survival of *Tcrδ*^+*/*+^*Ggta1*^*-/-*^ (n=7) and *Tcrδ*^*-/-*^*Ggta1*^*-/-*^ (n=8) mice infected as in (A); 2 experiments. **H**) CFU of Ae and An bacteria of *Tcrδ*^+*/*+^*Ggta1*^*-/-*^ (n=5) and *Tcrδ*^*-/-*^*Ggta1*^*-/-*^ (n=5) mice, 24 hours after infection; 2 experiments. Symbols in (B, D, F, H) are individual mice. Red bars in (B, D, F, H) are median values. P values in (A, C, E, G) calculated using log-rank test and in (B, D, F, H) using Mann-Whitney test. Peritoneal cavity (PC). *P < 0.05; ns: not significant

**Figure S3. The protective effect of IgG NAb acts irrespectively of αGal recognition. Related to Figure 3.**

**A**) Concentration of IgG in serum from *Ggta1*^+*/*+^ (n=10) and *Ggta1*^*-/-*^ (n=10) mice; 2 experiments. **B**) Representative single stained control plots for *Fig. 3D,E* showing *Rag2*^*-/-*^*Ggta1*^*-/-*^ cecal bacteria stained with BSI-B4 lectin for αGal, purified IgG from *Ggta1*^+*/*+^ and purified IgG from *Ggta1*^*-/-*^ mice. **C**) Representative flow cytometry plots showing IgG-binding of *in vitro*-grown species of bacteria isolated from the mouse microbiota incubated with IgG purified from *Ggta1*^+*/*+^ or *Ggta1*^*-/-*^ mice, as indicated in *Fig. 3F*. **D**) Representative flow cytometry plots of αGal expression by *in vitro*-grown species of bacteria isolated from the mouse microbiota, as indicated in *Fig. 3G*. Plots for *E. faecalis* (C, D) are highlighted in blue. Symbols in (A) are individual mice. Red bars in (A) are mean values. Error bars (A) correspond to SD. P values in (A) calculated using Mann-Whitney test. ns: not significant

**Figure S4. IgG from *Ggta1*^+*/*+^ and *Ggta1*^*-/-*^ mice recognize cecal bacteria to the same extent. Related to Figure 4.**

**A**) Median Fluorescence Intensity (MFI) of IgG^+^ bacteria from the same samples as in *Fig. 4A*. **B**) MFI of IgG^+^ bacteria from the same samples as in *Fig. 4B*. **C-F**) Representative controls plots for data shown in *Fig. 4C,D* showing *Rag2*^*-/-*^*Ggta1*^*-/-*^ cecal bacteria stained with **C**) PECy5-conjugated *Ggta1*^+*/*+^ IgG co-stained with AF594-conjugated *Ggta1*^*-/-*^ IgG, **D**) PECy5-conjugated *Ggta1*^*-/-*^ IgG co-stained with AF594-conjugated *Ggta1*^+*/*+^ IgG, **E**) PECy5-conjugated *Ggta1*^+*/*+^ IgG co-stained with AF594-conjugated *Ggta1*^+*/*+^ IgG and **F**) PECy5-conjugated *Ggta1*^*-/-*^ IgG co-stained with AF594-conjugated *Ggta1*^*-/-*^ IgG. **G-H**) Relative abundance of **G**) IgG^+^ bacteria and **H**) IgG^-^ bacteria at >2% frequency in the same samples as in *Fig. 4E-H*. Stacked bars represent the mean and colors represent the relative fraction of each taxon. Symbols (A, B) are individual mice. Red lines (A, B) are mean values. Error bars (A, B) correspond to SD. P values in (A, B) calculated using Mann-Whitney test, ns: not significant.

**Figure S5. Controls for *in-vivo* phagocytosis assay, detection of αGal on purified IgG and IgG binding to other recombinant mouse FcγR. Related to Figure 5.**

**A**) Analysis of peritoneal neutrophils based on the expression of CD11b and Ly6G in the same samples as in *Fig. 5A-C*. **B**) Bacterial uptake by peritoneal CD45^+^ leukocytes recovered from mice detailed in (A). **C**) Analysis of phenotype of phagocytic CD45^+^ cells from (B). **D**) Control of *Fig. 5D,E* for detection of αGal with BSI-B4 lectin in BSA conjugated to αGal and unconjugated BSA. SDS gel is shown as loading control. **E-H**) Relative binding to mouse **E**) FcγRI, **F**) FcγRIIb, **G**) FcγRIII and **H**) FcRn by *Ggta1*^+*/*+^ (n=4-7) and *Ggta1*^*-/-*^ (n=3-6) purified IgG, where n corresponds to independent IgG preparations. Data is representative of 1-3 independent experiments. Error bars correspond to SEM. P values calculated using 2-Way ANOVA with Sidak’s multiple comparison test.

**Figure S6. Epidemiological contexts where survival advantage against infection outweighs the reproductive fitness cost associated with loss of *Ggta1*. Related to Figure 6.**

Contour plots showing fitness ratios of *Ggta1*^*-/-*^ and *Ggta1*^+*/*+^ genotypes. Lifetime exposure to infection (*E*), assumed as constant over age, is plotted on the x-axis. The magnitude of protection (*p*) is plotted on the y-axis. Black line indicates the threshold for positive selective advantage (*s>0*) of *Ggta1*^*-/-*^ *vs. Ggta1*^+*/*+^ genotype. **A**) Condition of low virulence (*v*) relative to natural mortality. The selective forces for protection are lower than the cost of reproduction for the *Ggta1*^*-/-*^ genotype. **B**) Condition of higher virulence relative to natural mortality. Here, a parameter region above the black line emerges where the survival advantage exceeds the cost. **C**) Condition of further increase in virulence, increasing the possibility of selection even for lower range of exposure and protection. **D**) Condition of very high virulence, which favors selection for an even smaller protective effect.

## Notes

### Competing Interest Statement

The authors have declared no competing interest.

