## Supplementary figures and images for "A trade-off between resistance to infection and reproduction in primate evolution"

### Supplemental Figure 1

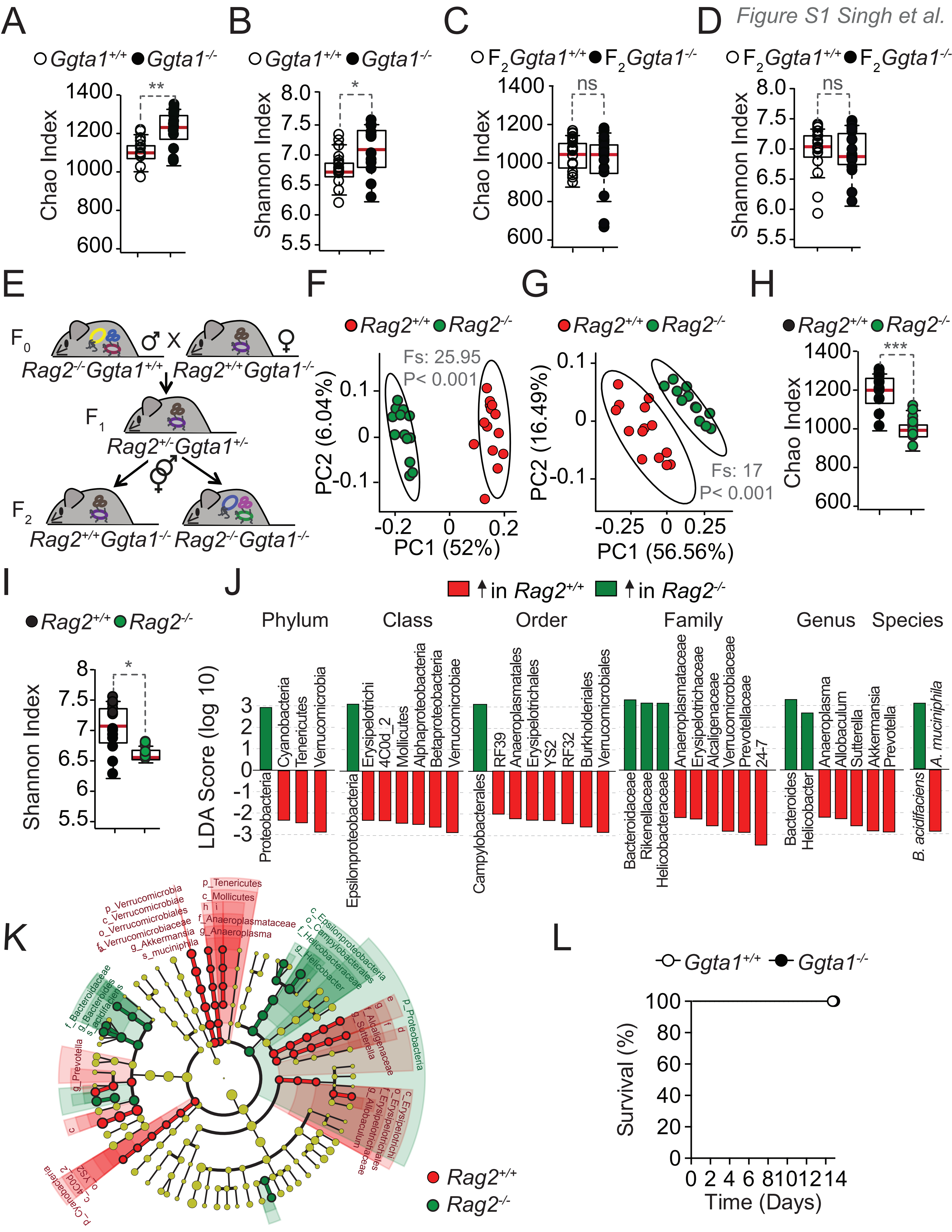

### Supplemental Figure 2

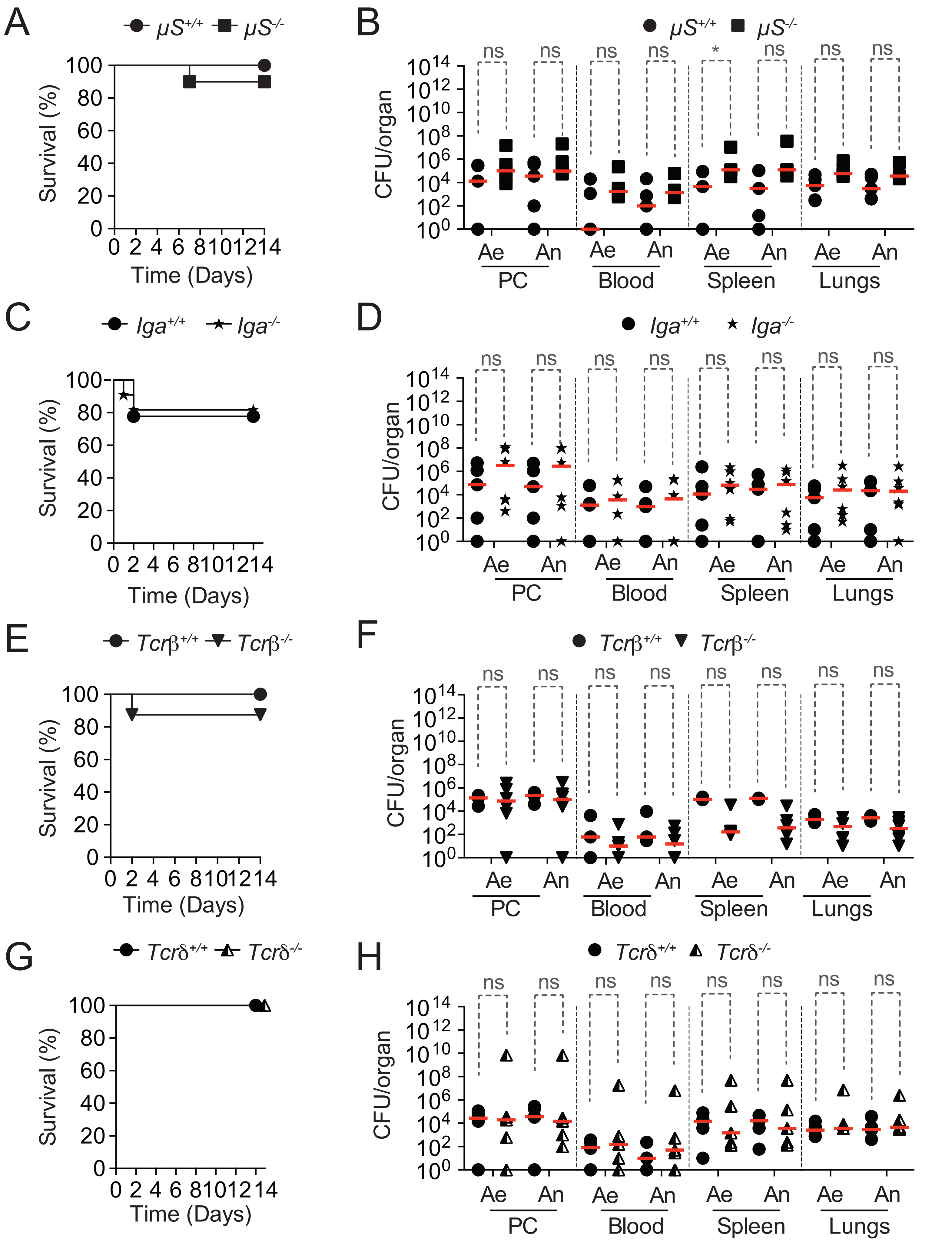

### Supplemental Figure 3

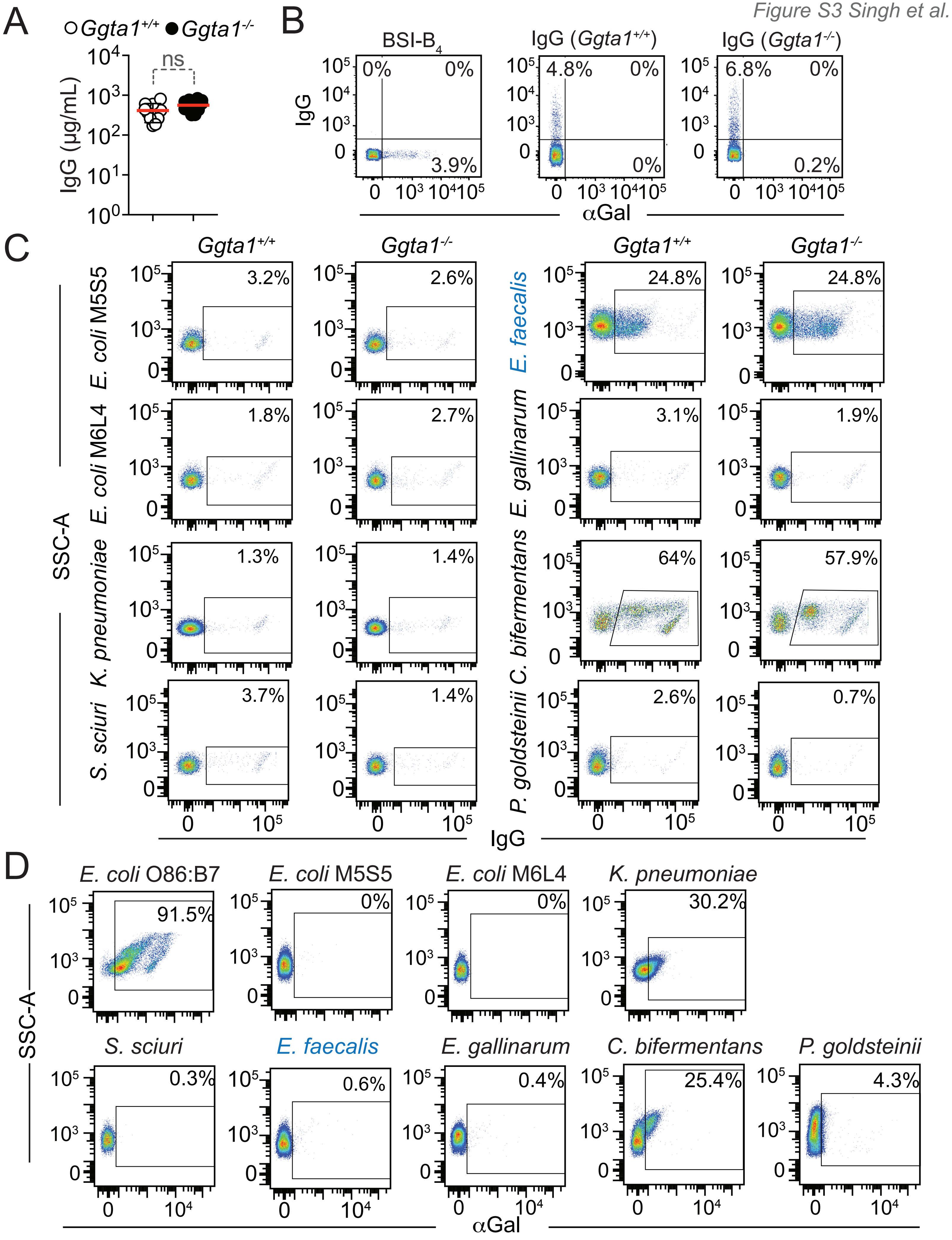

### Supplemental Figure 4

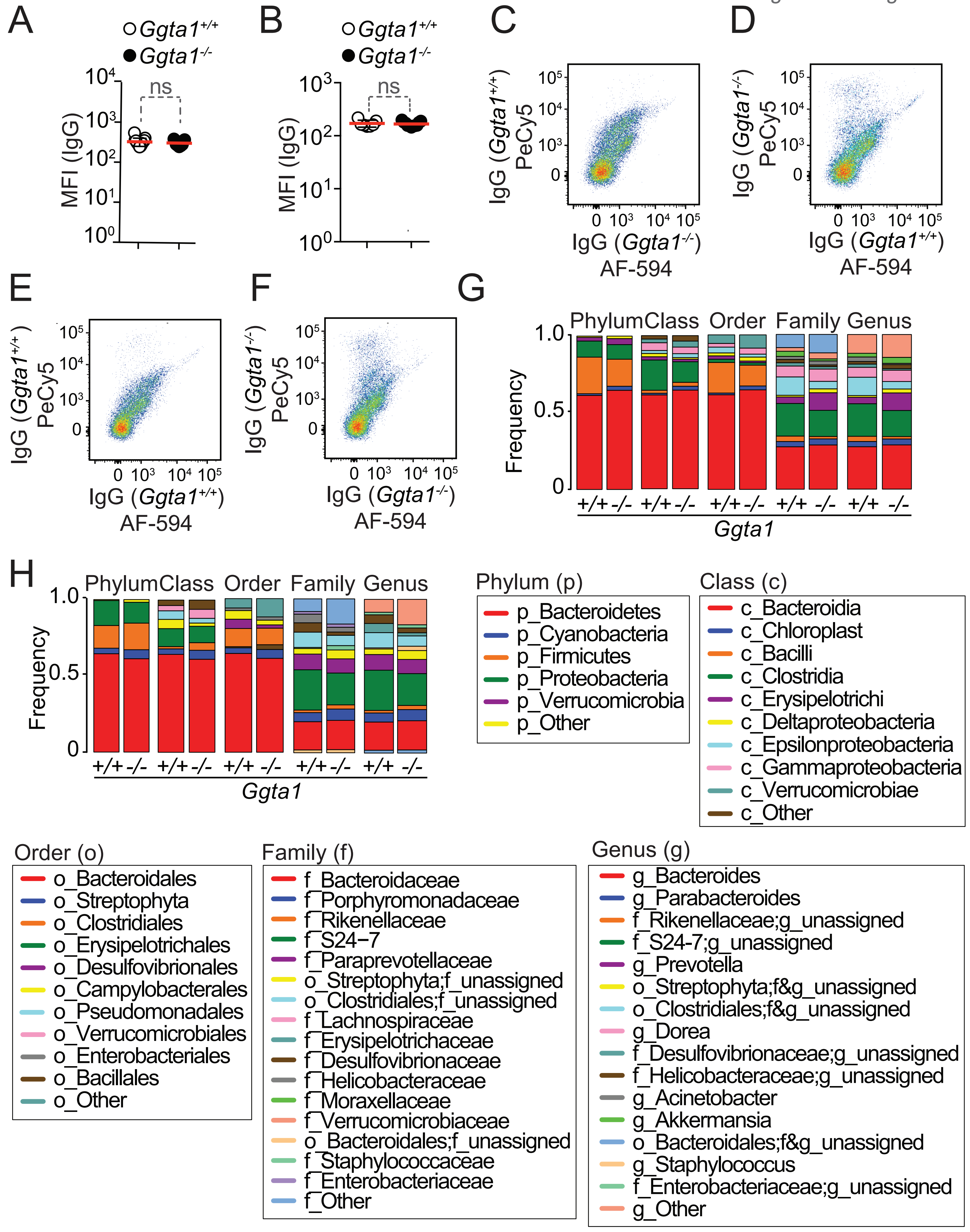

### Supplemental Figure 5

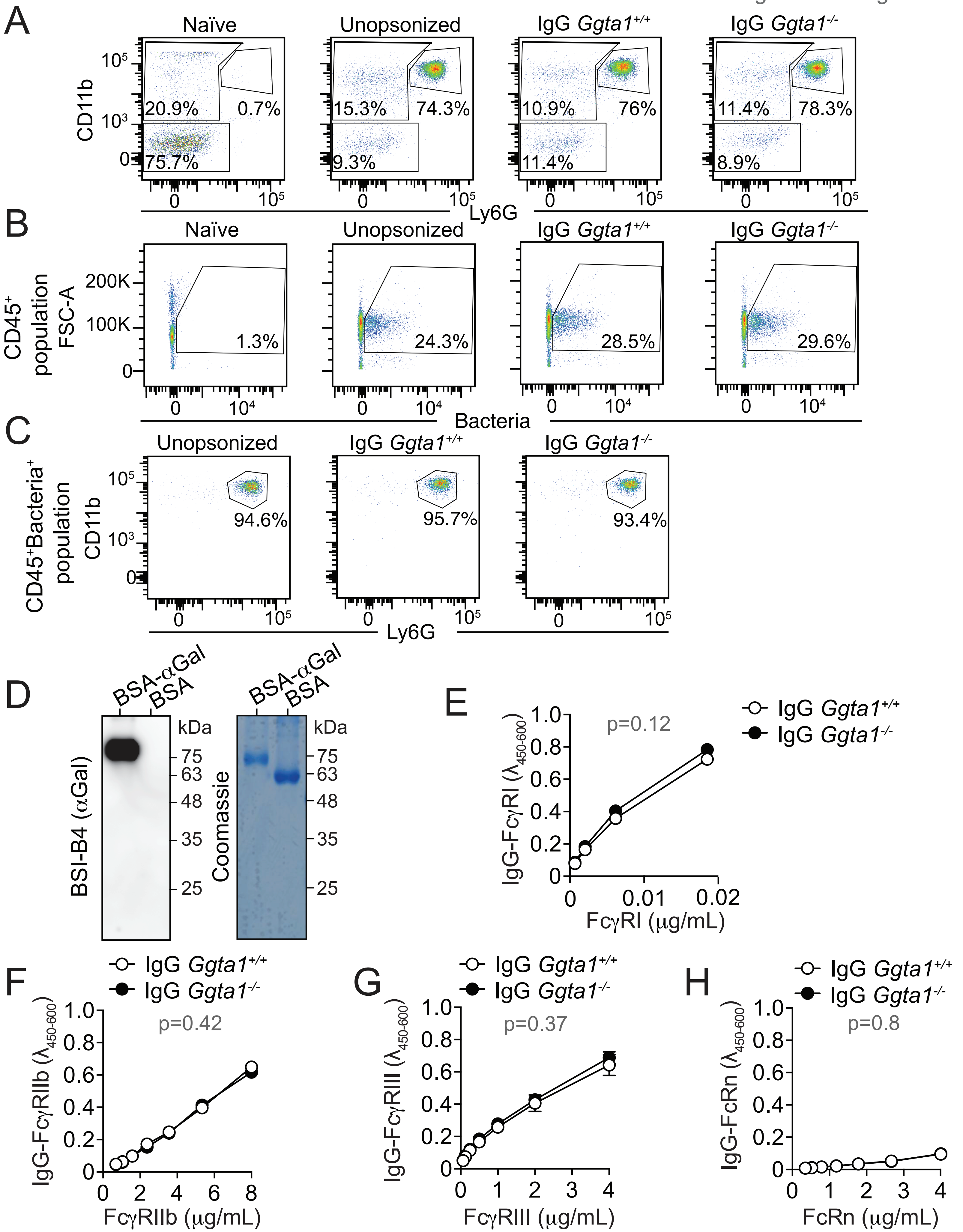

### Supplemental Figure 6

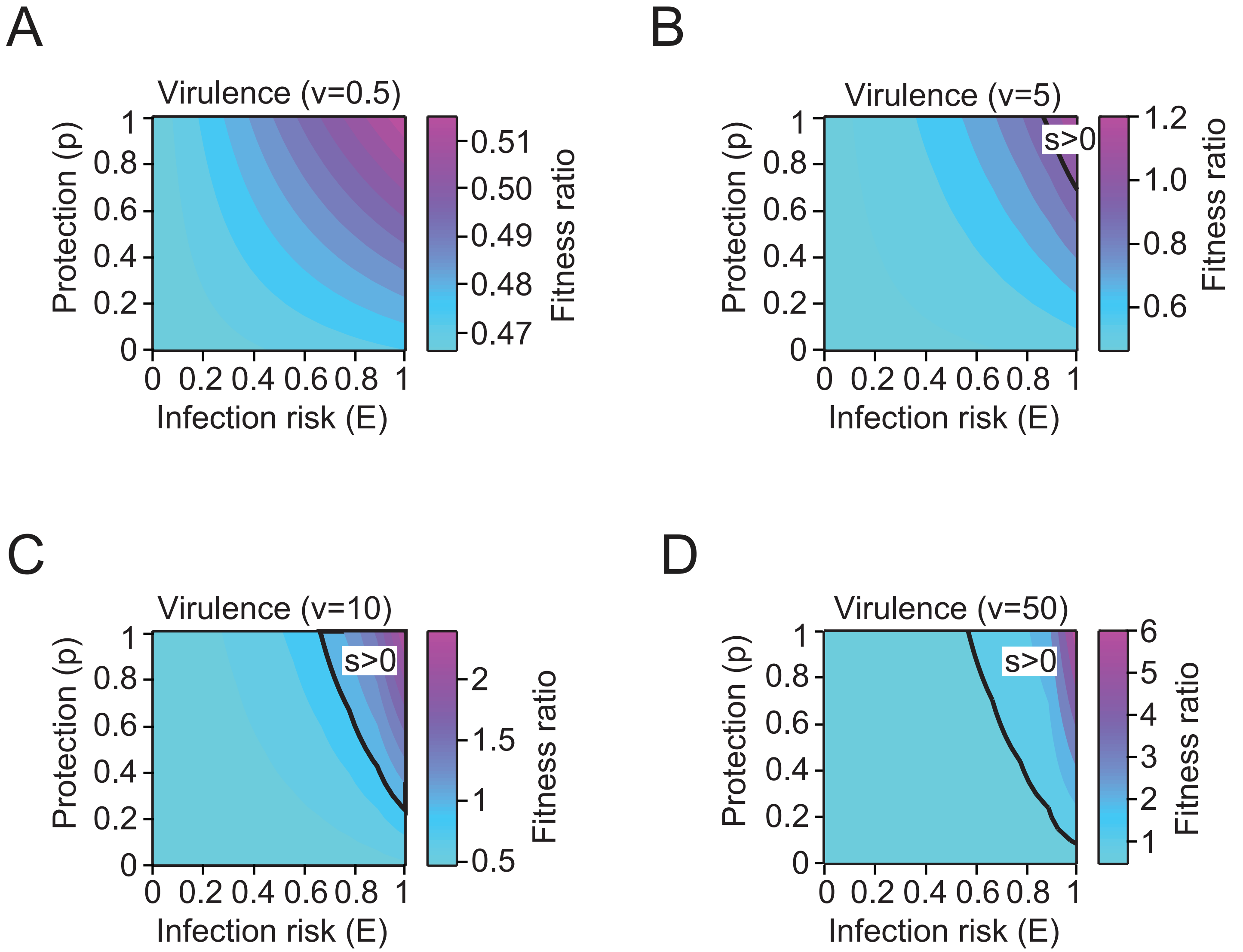
